# Can associative learning be the general process for intelligent behavior in non-human animals?

**DOI:** 10.1101/2021.12.15.472737

**Authors:** Johan Lind, Vera Vinken

## Abstract

The general process- and adaptive specialization hypotheses represent two contrasting explanations for understanding intelligence in non-human animals. The general process hypothesis proposes that associative learning underlies all learning, whereas the adaptive specialization hypothesis suggests additional distinct learning processes required for intelligent behavior. Here, we use a selection of experimental paradigms commonly used in comparative cognition to explore these hypotheses. We tested if a novel computational model of associative learning — A-learning — could solve the problems presented in these tests. Results show that this formulation of associative learning suffices as a mechanism for general animal intelligence, without the need for adaptive specialization, as long as adequate motor- and perceptual systems are there to support learning. In one of the tests, however, the addition of a short-term trace memory was required for A-learning to solve that particular task. We further provide a case study showcasing the flexibility, and lack thereof, of associative learning, when looking into potential learning of self-control and the development of behavior sequences. From these simulations we conclude that the challenges do not so much involve the complexity of a learning mechanism, but instead lie in the development of motor- and perceptual systems, and internal factors that motivate agents to explore environments with some precision, characteristics of animals that have been fine-tuned by evolution for million of years.

**Author summary:** What causes animal intelligence? One hypothesis is that, among vertebrates, intelligence relies upon the same general processes for both memory and learning. A contrasting hypothesis states that important aspects of animal intelligence come from species- and problem specific cognitive adaptations. Here, we use a recently formulated model of associative learning and subject it, through computer simulations, to a battery of tests designed to probe cognitive abilities in animals. Our computer simulations show that this associative learning model can account well for how animals learn these various tasks. We conclude that a major challenge in understanding animal and machine intelligence lies in describing behavior systems. Specifically, how motor flexibility and perceptual systems together with internal factors allow animals and machines to navigate the world. As a consequence of our results, together with current progress in both animal- and machine learning, we cannot reject the idea that associative learning provides a general process for animal intelligence.

## Introduction

Where does animal intelligence come from? This longstanding question has been approached by many and has given rise to a separation between ideas about a general process and ideas regarding adaptive specialization [1]. Within this central dichotomy, the general process approach proposes that associative learning underlies all learning. In contrast, adaptive specialization suggests there are additional learning processes, distinct from associative learning, that support intelligent behavior.

More specifically, the general process hypothesis posits that intelligence stems from a few general learning and memory mechanisms. These mechanisms are alike in different species and enable animals to solve a wide variety of problems. Observations of similar learning and memory functions among vertebrates have been brought up to support this view [1–4]. For example, the observation that regardless of species, the same training methods based on general concepts of learning can be used to train animals successfully on a diversity of tasks [6–8]. Training principles are the same for bees, chimpanzees, pigeons, rats, ravens and dogs. Although learning is generally considered to be associative, the exact content of what exactly is learned is still up for debate [9–11]. While the general process view argues for universal learning and memory mechanisms, it does not exclude species differences. Variation is explained in terms of genetically guided learning. Animals are genetically predisposed to learn/attend to specific stimuli and/or responses. This has also been referred to as ‘constraints of learning’ or ‘genetic predispositions’ [12, 13]. The general process approach has been criticized for not acknowledging the evolutionary history of species under investigation, which is important in order to compare species and to expand findings from one species to another [14].

The literature on adaptive specialization, instead, argues that intelligence arises from species specific mechanisms that evolved through natural selection in response to niche specific problems [15]. The idea of general process is replaced by species unique collections of computational mechanisms for learning and problem solving. This ecological view on intelligence is supported by, for example, observations of specialized behaviors that fit a species niche. Examples of this are rats possessing a unique type of learning about olfactory cues [16, 17], some species processing quantities of stimuli (numerosity) [18], or that some species, through convergent evolution, have evolved planning capacities [19]. It is argued that general learning mechanisms are insufficient for explaining these distinct capacities, which suggests that species evolve specific mechanisms for solving specific problems. Due to the species specific nature of the adaptive specialization framework, this hypothesis does not yet allow for an overarching theory on how different learning mechanisms work, so that the understanding of, for example, olfactory learning in rats cannot be extended to studies on numerosity. This has led to critique in that such ‘ecological’ accounts may be ultimately untestable [1].

Whether animal intelligence arises from general processes or adaptive specializations is still unresolved, as evidenced by a recent special issue in Frontiers in Psychology. Here, a total of 20 articles discussed general process versus adaptive specialization under the title “The Comparative Psychology of Intelligence: Macphail Revisited” [20]. One of the key issues raised here is the role of associations and associative learning in animal cognition. According to the general process view, animal intelligence depends at large upon associative learning. However, many researchers remain skeptic of the power of associative learning, particularly within comparative cognition and cognitive ethology where associative learning is regularly seen as too elementary to account for complex behaviors [19, 21, 22]. Researchers, therefore, suggest that animal intelligence does not arise from associative learning exclusively, but from additional components, such as “mental representation”, “rule-based learning” and “symbolic processing” [22], or for instance “sophisticated technical intelligence”, “complex problem solving” and “future-directed thought” [19]. No consensus was reached in this special issue, and the debate on general process versus adaptive specializations is still ongoing.

As a general understanding of intelligence in both AI and animal cognition is still out of reach, Matthew Crosby, Benjamin Beyret, and Marta Halina created The Animal-AI Olympics [23]. They pointed out limits in modern artificial intelligence models and argued that a radical change in research methodology is needed to better understand intelligence. They provide a testbed where agents are subjected to simple environments with objects of different kinds and rewards, all obeying physics rules (see http://animalaiolympics.com/AAI). For the testbed, they collated standardised and representative animal tests that together probe a general set of cognitive abilities, representing simplified and complex examples of natural challenges animals have evolved to solve [24].Submitted agents competed in a series of environments inspired by animal studies [24]. By combining AI and animal intelligence research, their focus on general aspects of intelligence based on comparative cognition research may provide a fruitful approach for synthetic work. In addition, their work highlights the need for theoretical approaches to understand animal intelligence [25–27].

Most entries that were submitted to The Animal-AI Olympics used Deep Reinforcement Learning algorithms [24]. The winner used a common three-step approach where first training environments were made, after which an agent received training, ending with a validation step involving behavior analysis. These Reinforcement Learning algorithms and recent models in animal learning research have much in common [28–30], and by taking both general process and adaptive specialization into account they may provide useful tools for working towards a synthesis of animal intelligence. By bridging the gap between comparative cognition and AI, The Animal-AI Olympics provide a new benchmark selection of tests different from other AI tests investigating intelligence, such as Arcade Learning Environment (ALE) [31], OpenAI Gym [32], and General Video Game AI [33]. We believe Crosby et al. provide a novel starting point for studies of intelligence [23, 24], by welcoming theoretical studies that can be compared with results from the empirically driven field of comparative cognition.

In this paper, we take advantage of the benchmark tests selected for The Animal-AI Olympics [23, 24] and subject a new associative learning model — A-learning — to these tests [30, 34]. This recently developed model can produce long behavioral sequences and can yield optimal solutions to problems encountered by an agent. Besides, it can account for general observations of social learning in animals [35] and planning behavior observed in great apes and ravens [26]. In addition, this model can reproduce core features of results from animal psychology, for example instrumental and Pavlovian acquisition, conditioned reinforcement, and different kinds of higher-order conditioning [30]. The general nature of the included benchmark tests can further our knowledge of what role associative learning can play for animal intelligence. We also aim to inform the long-lasting conflict between general process and adaptive specialization as explanations for animal intelligence.

To subject this new model to the different tests, we translated the tasks included in The Animal-AI Olympics into script-based description of the environments and their reward structures. As this model does not include perceptual and motor mechanisms, environments and agents’ behaviors are translated into verbal descriptions of key stimuli and actions, as in previous computational analyses of learning phenomena with this model [26, 30, 34, 35]. In computer simulations, the agent learns from actions it performs to navigate the world of stimuli and rewards. These learning simulations can inform us on what is possible to learn through associative learning and what associative learning can do for animal intelligence. Answering detailed questions about species specific performance in different tasks is beyond the scope of this study, although we will touch upon this matter briefly. We will show that this model of associative learning alone provides a general process that can account for learning in almost all these tasks. One task required an additional memory mechanism for associative learning to perform successfully. To explore general questions about inter-specific variation, we also performed simulations to highlight how genetic factors can affect learning and test performance, and thereby animal intelligence.

## Materials and methods

Here, we first briefly describe A-learning, the associative learning model used for these simulations, and we also introduce the simulation software used in this study. We give detailed accounts of all tasks that have been subjected to computer simulations and end with a description of a case study where we use one of these tasks to explore different parameter settings.

### Associative learning model

To explore what associative learning can achieve if subjected to this collection of tests, we used a new formulation of associative learning called *A-learning* [30, 34]. Here, we will describe the model briefly, for further details please see [30]. At its basis, this model assumes a subject has a behavior repertoire and behaviors can be used to respond to a world of detectable environmental states. A behavior will take the animal from one state to another. At each such state, every stimulus has a genetically fixed primary reinforcement value, written *u*. This value can be negative, neutral, or positive, and it guides learning by favoring behaviors that maximize its total value, in line with the assumption that this favors survival and reproduction. Generally, consuming food will have a positive value, whereas something inflicting pain or distress will have a negative value. More importantly, this allows expectations about the value of a state to develop, making goal-directed behavior and learning of behavior sequences possible. The model includes two learning processes and one decision making rule. It is assumed that animals learn after experiencing an event sequence with a behavior *b* in response to a stimulus *s*, leading to the next stimulus *s*′, formalized as:

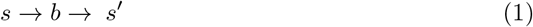

in which the model learns that performing behavior *b* to stimulus *s* yields a higher value if stimulus *s*′, is positive, than if *s*′, is neutral or negative. This value, written *v* (*s*→*b*), corresponds to the associative strength between stimulus *s* and behavior *b*. In functional terms this is stimulus-response (S-R) value learning, where *v* (*s*→*b*) represents an estimated value of performing behavior *b* when perceiving stimulus *s*. The second learning process, stimulus value learning, stores and updates the value of a stimulus. This estimates the value of *s*, which is updated according to the value of the subsequent stimulus *s*′, and written as *w*(*s*).

These two learning processes are integrated through the effect a stimulus value *w*(*s*) has on S-R value learning. The S-R value *v*(*s* → *b*) is updated according to the primary reinforcement value *u*(*s*′), the stimulus value *w*(*s*), and the previously stored S-R value according to the equation:

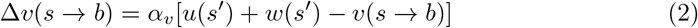

where Δ*v* (*s*→*b*) is the change in *v* (*s*→*b*) caused by the experience in the event sequence in Eq 1. In addition, *α*_*v*_ is a positive learning rate determining the rate of updating *v* (*s*→*b*) The stimulus value *w*(*s*) is updated according to the equation:

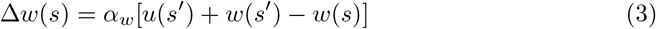

where *α*_*w*_ is a positive learning rate for the update of the stimulus value *w*(*s*). The stimulus value *w*(*s*) affects learning about behaviors and acts as a conditioned reinforcement, or secondary reinforcer. This allows for stimuli lacking an inborn primary reinforcement value to acquire stimulus value. In the above case, if *s*′ is positive, an animal experiencing the event triplet in Eq 1 will also attribute a positive value to experiencing *s*, which subsequently reinforces behavior that leads to experiencing *s*. At decision making a behavior is selected from the behavior repertoire. A-learning uses the softmax decision rule (see [30] for details). The probability of choosing *b* in response to *s* is then formally described as:

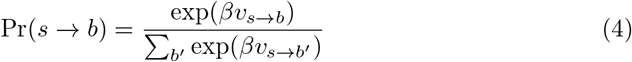

which includes the parameter *β* that regulates the amount of exploration. With *β*= 0, all behaviors are equally likely to be selected, meaning that prior experiences play no role in decision making. As *β* increases behaviors with the highest estimated value *v* will have a higher probability of being selected. This decision making process has been shown to match empirical observations [30, 34].

Let us use a practical example from one of the tasks included in The Animal-AI Olympics to exemplify A-learning, and how it can be described in line with Eq 1. For an animal to learn to make a detour it must learn to avoid a barrier. This behavioral sequence can be described in terms of its key events as:

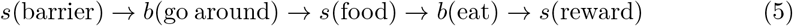

where the animal needs to experience the end of the sequence first to learn how to get food. Initially, only eating in the presence of food is rewarding, so an animal that starts from the beginning of the sequence must first perform initially non-rewarding behaviors, such as walking around the barrier to find the food and eat it, to learn the whole sequence. Let *s*(food) have an initial value of zero, meaning it is a neutral stimulus that does not act as a reinforcer. Eating the food results in experiencing *s*(reward) that has positive primary reinforcement value, thereby the animal learns to eat when subjected to *s*(food). In addition, the positive value of *s*(reward) makes the stimulus value of the preceding *s*(food) increase. This changes the situation for the animal, because if it chooses to go around the barrier it will again experience the stimulus *s*(food), but it has now acquired positive stimulus value. This way, *s*(food) acts as a reinforcer, making walking around the barrier rewarding in its own right, and the key aspect of this task can be learned (see table 1 for detailed descriptions of all tasks).

**Table 1.** Summary of the key aspects of the included learning scenarios.

| Task | What is learned? | Experience | Outcome |
| --- | --- | --- | --- |
| *Mazes* | Enter correct arm | $S_{\text{start}} \rightarrow B_{\text{go left}} \rightarrow S_{\text{food}}$ | $v(S_{\text{start}} \rightarrow B_{\text{go left}}) > 0$ |
| *Delayed gratification* | Wait in presence of food | $S_{\text{food}} \rightarrow B_{\text{wait}} \rightarrow S_{\text{more food}}$ | $v(S_{\text{food}} \rightarrow B_{\text{wait}}) > 0$ |
| *Detour* | Go around barrier | $S_{\text{barrier}} \rightarrow B_{\text{go around}} \rightarrow S_{\text{food}}$ | $v(S_{\text{barrier}} \rightarrow B_{\text{go around}}) > 0$ |
| *Cylinder task* * | Ignore blocked food | $S_{\text{food cylinder}} \rightarrow B_{\text{bump}} \rightarrow S_{\text{no food}}$ | $v(S_{\text{food cylinder}} \rightarrow B_{\text{bump}}) \downarrow$ |
| *Puzzle box* | 3 behaviors unlock door** |  |  |
| 1 | Claw string | $S_{\text{string}} \rightarrow B_{\text{claw}} \rightarrow S_{\text{food}}$ | $v(S_{\text{string}} \rightarrow B_{\text{claw}}) > 0$ |
| 2 | Press platform | $S_{\text{platform}} \rightarrow B_{\text{press}} \rightarrow S_{\text{food}}$ | $v(S_{\text{platform}} \rightarrow B_{\text{press}}) > 0$ |
| 3 | Lift bar | $S_{\text{bar}} \rightarrow B_{\text{lift}} \rightarrow S_{\text{food}}$ | $v(S_{\text{bar}} \rightarrow B_{\text{lift}}) > 0$ |
| *Spatial elimination* | Find food under board | $S_{\text{inclined}} \rightarrow B_{\text{take inclined}} \rightarrow S_{\text{food}}$ | $v(S_{\text{inclined board}} \rightarrow B_{\text{take}}) > 0$ |
| *Gravity bias* | 2 behaviors are learned |  |  |
| 1 | Food falls straight down | $S_{\text{food X}} \rightarrow B_{\text{take at X}} \rightarrow S_{\text{food}}$ | $v(S_{\text{food X}} \rightarrow B_{\text{take below X}}) > 0$ |
| 2 | Food follows other route | $S_{\text{food X}} \rightarrow B_{\text{take at Y}} \rightarrow S_{\text{food}}$ | $v(S_{\text{food X}} \rightarrow B_{\text{take at Y}}) > 0$ |
| *Radial maze* *** | Find all food in maze |  |  |
| 1 | Do not return | $S_{\text{center}} \rightarrow B_{\text{return}} \rightarrow S_{\text{little food}}$ | $v(S_{\text{center}} \rightarrow B_{\text{return}}) > 0$ |
| 2 | Do not go straight | $S_{\text{center}} \rightarrow B_{\text{straight}} \rightarrow S_{\text{little food}}$ | $v(S_{\text{center}} \rightarrow B_{\text{straight}}) > 0$ |
| 3 | Go even number of steps | $S_{\text{center}} \rightarrow B_{\text{go even}} \rightarrow S_{\text{more food}}$ | $v(S_{\text{center}} \rightarrow B_{\text{go even}}) > 0$ |
| 4 | Go odd number of steps | $S_{\text{center}} \rightarrow B_{\text{go uneven}} \rightarrow S_{\text{most food}}$ | $v(S_{\text{center}} \rightarrow B_{\text{go uneven}}) > 0$ |
| *Object permanence* | Learn to take correct cup |  |  |
| 1 | Food left, take left | $S_{\text{food left}} \rightarrow B_{\text{take left}} \rightarrow S_{\text{food}}$ | $v(S_{\text{food left}} \rightarrow B_{\text{take left}}) > 0$ |
| 2 | Food center, take center | $S_{\text{food center}} \rightarrow B_{\text{take center}} \rightarrow S_{\text{food}}$ | $v(S_{\text{food center}} \rightarrow B_{\text{take center}}) > 0$ |
| 3 | Food right, take right | $S_{\text{food right}} \rightarrow B_{\text{take right}} \rightarrow S_{\text{food}}$ | $v(S_{\text{food right}} \rightarrow B_{\text{take right}}) > 0$ |
| *Numerosity* | Take bowl with most food | $S_{\text{more/less}} \rightarrow B_{\text{take more}} \rightarrow S_{\text{more food}}$ | $v(S_{\text{more food}} \rightarrow B_{\text{take more}}) > 0$ |
| *Tool use* | Take cloth with food | $S_{\text{cloth food}} \rightarrow B_{\text{take cloth}} \rightarrow S_{\text{food}}$ | $v(S_{\text{cloth food}} \rightarrow B_{\text{take cloth}}) > 0$ |

\*The food rewarding behavior is learned in pre-training, in tests, avoiding to bump into cylinder is recorded. \*\*Food can only be collected after all three behaviors, in any order, have been performed. \*\*\* Due to systematic differences in outcomes between behaviors, this will result in $v(S_{\text{center}} \rightarrow B_{\text{return}})$ and $v(S_{\text{center}} \rightarrow B_{\text{straight}}) < v(S_{\text{center}} \rightarrow B_{\text{go even}}) < v(S_{\text{center}} \rightarrow B_{\text{go uneven}})$ .
\*

### Learning simulator

To perform the simulations we used the newly developed Learning Simulator [36], an open source software designed for simulating learning in animals and humans. The simulator is publicly available at https://www.learningsimulator.org/ where documentation and some example scripts can be found. The Learning Simulator has previously been used to explore planning behavior [26], social learning [35], and general phenomena in experimental psychology [30]. The Learning Simulator frames learning using a subject that interacts with an environment in accordance with an event sequence as described in Eq 1. A subject perceives stimuli from the environment to which it can respond with some behavior, which in turn makes the environment present the next stimulus to the subject. The Learning Simulator can be used with several different animal- and reinforcement learning algorithms, but here we have used A-learning throughout [30, 34]. With the exception of the object permanence task we used the A-learning mechanisms alone throughout. There is no discriminative stimulus present in the choice situation of the object permanence task, making this task impossible for A-learning to solve without some kind of memory of the preceding stimulus (see description in Method section). It is well known that animals can remember arbitrary stimuli for short times through memory traces [37, 38], a short-term memory that is an integrated part of animals’ general processes of learning and memory [39, 40]. To explore if associative learning with an integrated trace memory could solve the object permanence task, we used a feature of the Learning Simulator that integrates a trace memory with A-learning. This way, a preceding stimuli can be used for decision making in the present, at exponentially decaying intensities *θ*, where *θ* < 1. For this task we used the integrated trace memory with *θ* = 0.6.

### Simulation details

The below scripts are available as text-files at the following url: https://doi.org/10.17045/sthlmuni.17068409. These files can be used in the above described Learning Simulator to perform each respective computer simulation. All computer simulations are based on the studies selected for the benchmark tests in The Animal-AI Testbed (http://animalaiolympics.com/AAI/testbed). The order of the tasks is the same as presented there. Based on the descriptions of the tasks in the literature, we identified the key events that are critical for learning each task and included these in scripts. We focused on the stimuli, objects and other components in the environment that are of importance for the animal to perform in the task. All experiments end with some reward. Besides, we looked at what behaviors the animal could respond with towards the stimuli available and what influence these responses have on the environment. In some cases, these key events relied on behaviors that were trained before the start of the experiment (pre-training), for example specific responses to objects. If the test results depended on choices made and behaviors learned during pre-training, these are included in our scripts.

In order to make the simulations realistically comparable to what animals experience when they enter the test, we assumed that the animals in our simulations possess general everyday motor and perceptual skills. In terms of motor skills we assumed that the animals are able to move between parts of an experimental area, reach for or explore hidden objects that may be inside some contraption or behind a wall, and to identify and consume food rewards. In terms of perception, we assumed that stimuli could be identified and if overtly distinct could be told apart. Assumptions about perception and motor skills means that these specific abilities did not have to be learned prior to a task. Instead, the simulations focus on the behavior sequence unique to the experiment at hand, adhering to the logical order of stimulus presentation, reward structure, and behaviors performed.

In addition to task specific behavior we always included the possibility for a subject to ignore any stimulus, equivalent to not reacting. Apart from in the final case study on variation (see section A case study on variation in learning and test performance), we used the same learning parameters in all tasks to facilitate comparisons between tasks. Learning parameters were set to: learning rate *α* = 0.2 for updates of both stimulus-response- and stimulus values, exploration was set at *β* = 0.5, behavior cost=0.1, and rewards *u* = 10. Simulations were performed with 500 subjects. Any departure from these setting will be highlighted in the text. We would also like to mention that replications of these simulations will result in similar but not necessarily identical results due to the stochastic nature of the decision-making equation.

Here follows detailed descriptions of all tasks included and the sources that inspired the tasks in the Animal-AI Olympics. All key experiences and learning outcomes are presented in table 1.

#### Two-choice mazes (Y- and T-maze)

*Descriptions from Crosby et al*. [24]: “A Maze in the shape of a Y. One branch contains a preferred reward to the other and usually both can be seen at the same time.” And a T-maze is describes in similar terms, with the key difference that in a T-maze a reward cannot be perceived from the starting point: “Like a Y-Maze except that both arms are not visible at the same time.” Here, we simulated a case where the animal starts in one arm (the ‘start’ arm) and can either make a right or a left turn into the two other arms of the maze. Given the fact that these behaviors would look the same for both the Y and the T-maze, we have grouped these together for our simulations. Either the left or the right arm contains a food reward, as in for example a study on cows [41] and a study on nematodes [42]. To accomplish this we use a ‘start stimulus’ to which the behaviors ‘go to the left’ and ‘go to the right’ are possible. Choosing go to the left results in an opportunity to eat a food reward. (Filename of script: 1-Mazes.txt.)

#### Delayed gratification

*Description from Crosby et al. 2020* [24]: “The ability to forgo an immediate, less preferred reward for a future, more preferred reward. Solving this robustly in the Animal-AI environment requires understanding that there will be a larger delayed reward based on the physics of the environment.”

We simulated a case where great apes were presented with a bowl that was within reach, and an experimenter placed pieces of food into the bowl one by one [43]. If the ape pulled the bowl into the cage the trial ended and no more food was given that trial. In a situation where rewards accumulate as long as they are not collected, an animal would gain more rewards by waiting instead of immediately collecting a small reward. In this script, we made it possible to collect more than one food item by including three behaviors: ‘take the food’ (meaning that no more food will be collected), and ‘wait’, and ‘eat the food’ (only possible once food has been collected). (Filename of script: 2a-Delayed-gratification.txt.).

#### Detour task

*Description from Crosby et al. 2020* [24]: “Testing the ability to make a detour around an object to get food and assess the shortest path to the object.” Here we simulated a case inspired by a study on dingoes that were presented with a V-shaped mesh fence [44]. They could see a food reward through the fence, only being able to obtain it by making a detour around the outside of the fence. Going straight towards the fence would not lead to the dingo receiving the food reward. To get to the food, the dingoes had to walk away from the food, reach the end of the fence and then turn around to walk back towards the food. Detour tasks are similar to delayed gratification tasks and self-control studies in that behaviors such as ignoring, waiting, or walking away from food needs to be learned. In detour tasks, in addition to ignoring something, an animal also needs to learn to go around some kind of barrier. In this simulation, going towards the barrier instead of around it needs to be overcome. Behaviors included are getting stuck at the barrier, going around the barrier from two positions (start position and position at the barrier, respectively) and ignore. To capture the difficulty of the task, namely that an animal is initially more likely to go straight towards the food than around the barrier, simulations began with a positive value for going to barrier where food is visible. In addition, to make going around the barrier less likely simulations began with a negative value for going around the barrier to capture the spontaneous response to be attracted to the food, although it is initially out of reach. (Filename of script: 3-Detour.txt.)

#### Cylinder tasks

*Description from Crosby et al. 2020* [24]: “In the testbed this includes both opaque and transparent cylinders.” Just like in detour tasks and delayed gratification tasks, to solve cylinder tasks an animal must learn to overcome reaching for a food item that is visible through a transparent cylinder but not possible to grasp directly. We simulated a well known case where an animal first learns to find a piece of food inside an opaque cylinder. Afterwards the animal is subjected to a transparent cylinder with a now visible piece of food in the cylinder. To successfully retrieve the food the animal needs to perform the same behavior as in the opaque phase, and inhibit to try to grasp the food through the transparent cylinder, thereby bumping into the cylinder. The key for an animal to solve this task is to learn the correct behavior, that is reaching for the food from the side at the opening of the cylinder. The simulation follows the procedure of cylinder tasks with a training phase in which the food collecting behavior is learned, and a second phase when it is tested if, and how quickly, the apparent and incorrect behavior is inhibited. We included the incorrect bump the cylinder-behavior and the food collecting detour response. This situation described a behavior sequence so we let the detour response produce a food item that subsequently needed to be eaten to be rewarding. To capture the difficulty of the cylinder task we used the two phases. The simulation followed the procedure in that first a training phase occurred, whereby the correct behavior was learned to get the reward. Then followed a test phase that simulated the visible food and that an incorrect response towards the food needed to be inhibited to get the food. This was simulated by adding the stimulus food inside the cylinder in the test phase and a response towards that food based on the assumption that the animal recognizes the food and upon its presentation reaches for the food. The value of incorrectly reaching the food was set equal to the primary value of the food. (Filename of script: 4-Cylinder-tasks.txt.)

#### Thorndike’s puzzle box

*Description from Crosby et al. 2020* [24]: “These are recreations (of different realism) of Thorndike’s experiments on cats, dogs, and chicks where the agent must escape from a confined area and food is placed outside.”

Here we simulated one of the classic puzzle boxes designed by Thorndike for his thesis work [45]. An animal is confined in a box and it can ‘escape’ the box and collect a food reward outside by performing a set of specific behaviors. We used one of the more complex puzzle boxes (Box ‘k’) that required the animal to perform three separate actions to open the door. The door would only open after all three behaviors were performed, that is pulling/clawing a wire, press a platform on the floor, and lift/push down a bar on the door. The order in which the behaviors are performed does not matter. This introduces the problem that correct responses, such as lifting one of the bars, can be made without having a direct positive consequence. The animal needs to keep exploring and perform all three actions in order to associate any of these actions with the reward. To capture the large amount of possible actions the animal can explore once inside the box, the following behaviors are included in the script: lift, claw, press, go out, ignore, approach. Pressing and clawing behaviors are possible to all stimuli, lifting is not possible for the walls, the floor and the door. (Filename of script: 5-Thorndike-puzzle-box.txt.)

#### Spatial elimination

*Description from Crosby et al. 2020* [24]: “Spatial properties can be used to infer the location of food. For example, it can not be in the open space so if there is any it must be behind that wall.” To simulate this task we used a case where great apes were presented with two plastic boards. One plastic board was inclined because a food reward was hidden underneath. They were rewarded if touching the inclined board that hid food. (The test called Shape, see supplementary information in [46].) This translates to a situation where an animal needs to choose between two different stimuli, where choosing one stimulus results in food. Logically, this test resembles a two-choice maze. We included the following behaviors: take an inclined board, take a flat board and eat the reward (and ignore). If the inclined board was taken, a food reward could be collected, if the flat board was chosen a new trial started without any reward. (Filename of script: 7-Spatial-elimination.txt.)

#### Support and gravity bias

*Description from Crosby et al. 2020* [24]: “Tasks involving gravity and food supported on other objects.” Here, our simulations aim to explore the crux of these experiments, namely that animals expect objects to fall straight down if they are used to them doing so, even in novel situations. Crosby et al. (2020) use a study by [47] which involves cotton top tamarins performing a gravity bias task. Since the study at hand here is a follow up to the original gravity bias studies done in [48], we have used the original description for our simulations. Therefore, we mimicked the study by [48] that involved an apparatus consisting of two levels. The upper level has three openings that can be connected to three food boxes on a lower level. Food can be dropped through the openings in the top and end up in one of the boxes on the bottom, depending on the configuration of the tubes. This way, a food item dropped in the tube opening on the far left side may end up in the right container on the bottom, and so on. These experiments aimed to test if cotton top tamarins behave as if they expect food to fall straight down, which would be expected if they have a gravity bias. Contrary to this, the tamarins were expected to make the correct choice, and obtain the food, if they would use information about the bends and connections of the tubes. To simulate this in a faithful way, we designed a script including 3 phases. The first phase is constructed so the subjects learn about gravity. This is comparable to animals in reality, experiencing objects in their surroundings falling down. Rewards fall from openings in the upper level straight down to the bottom level. No Tubes are connected in this phase. In the second phase, the subjects experience the same training as the animals in [48] and learn to collect food that is hidden behind doors. Phase 3 only contains one configuration of the tube, the same as in the test phase of the study, where the tube goes from opening 1 on the top of the apparatus to door 3 on the bottom. The stimuli used in this script are s(up1), s(up2) and s(up3) for the food being dropped from the top level openings. s(d1), s(d2), and s(d3) represent food being put directly behind the doors in the lower level locations. s(b1), s(b2), and s(b3) represent the actual boxes. Food can get there through the tube or by being put in there directly. The behaviors used are b(go1), b(go2), and b(go3), which represent the animal opening the lower level doors. (*Filename of script* : 8-Gravity-bias.txt)

#### Radial mazes

*Description from Crosby et al. 2020* [24]: “Mazes with a number of spokes radiating out from a central hub.” Here we simulated an animal navigating a radial maze with eight arms where one food item per arm can be collected in each trial [49]. During a trial, the consumed food is not replaced. This way, the task tests if the animal learned to avoid already visited and thereby depleted arms. This simulation investigate two facets of the experiment. Firstly, if solving the task is possible for the subjects, and secondly whether a regular “algorithmic” behavior pattern emerges. This means that an individual would systematically perform just one out of several possible correct behaviors. As food items were hidden underneath cups in each arm, the maze looks the same irrespective of which arm the animal exits. Our script focuses on the behavioral options available and we therefore included the following behaviors: go one, two, or three steps to the left or to the right, go straight or return to the last visited arm. We also included a behavior to approach the cup where food was collected. Importantly, a food reward can only be collected upon the first instance an arm is visited, afterwards the arm is depleted. The stimuli used represent the start of the trial, the center of the maze and the inside of the arms with a cup under which the food is hidden. (Filename of script: 9a-radial-maze.txt and 9b-radial-maze-1ind.txt.)

#### Object permanence

*Description from Crosby et al. 2020* [24]: “These tests all involve food that moves out of sight that the agent needs to still attain.” We simulated the test called “Object permanence” [46] where great apes were presented with two or three empty cups on a platform. After presentation, an experimenter put a food reward under a smaller additional opaque cup. Subsequently, the smaller cup was moved underneath one of the larger cups. Now, the ape was allowed to choose one of the cups, and if the large cup with the smaller cup hidden underneath was chosen, the ape could collect the food reward. If one of the wrong cups was chosen the trial ended. We included the following behaviors: take left cup, take the right cup and take the center cup. However, an important aspect of this experiment is the disappearance of a stimulus. Therefore, when the choice is made the situation is identical for all conditions. In this case there is no information for A-learning available at the time of decision making it impossible for this mechanism to solve this task at a higher level than random choice. For this reason, we add trace memory to the learning mechanism, and this simulation therefore combines A-learning with a trace memory. That animals represent past stimuli – such as the location of the small cup – as a fading trace is well described in the literature [37, 38, 40]. Practically, this means that the location of the small cup is stored in memory and is available in time steps after it was perceived, but at a fading intensity. (Filename of script: 10-Object-permanence.txt.)

#### Numerosity

*Description from Crosby et al. 2020* [24]: “These tests all involve counting to navigate to the compartment with the most food.” To simulate a numerosity test we used the “Relative numbers” test [46] where great apes were presented with two dishes baited with different amounts of food rewards of equal size. The subject was allowed to chose one of the two dishes. We followed the same sequence as in that test with trials in the following order: 5:1, 6:3, 6:2, 6:4, 4:3, 3:2, 2:1, 4:1, 4:2, 5:2, 3:1 and 5:3. This was a two-choice task and the following behaviors were included: take the dish with fewer pieces of food or take the dish with more food. Thus, it was assumed that the great apes could identify that a dish with five or two pieces of food is different from a dish with one food item. In the first trial one dish contained one food reward and the other dish zero (1:0). (Filename of script: 11-Numerosity.txt.)

#### Tool use behavior

*Description from Crosby et al. 2020* [24]: “These test are based on the ability to use the pushable objects in the arena as makeshift tools to get food. They are the most complicated in the testbed and extend to the ability to perform simple causal reasoning about the outcome of actions.” We simulated the test in which cotton-top tamarins were subjected between a forced-choice between a cloth on which food was located vs. a cloth with food laying beside it (problem: “On” in [50]). If a tamarin pulled the cloth on which food was located it could collect the food reward. A tamarin pulling the cloth without food on top it received no food reward. This experiment represents a two-choice situation and to simulate the discrimination between choosing a cloth with and without food on top the following behaviors were included: take a cloth, ignore a cloth, and eat a reward. The simulation is based on experiencing either of two compound stimuli, s(cloth, food outside cloth) or s(cloth, food on top of cloth). Here, if ignoring one of the compound stimuli the subjects will experience the next compound stimulus. If choosing to take the cloth it can result in the opportunity to eat a reward if *s*(cloth, food on top of cloth) was taken, but no reward and the end of the trial followed a choice of s(cloth, food outside cloth). (Filename of script: 12-Tool-use.txt.) There are many known instances of more complicated tool use in animals, see Enquist et al. [34] (section 5.1.: “Chaining in nature”) for an associative learning account of the ontogeny of stone tool use in chimpanzees.

#### A case study on variation in learning and test performance

To supplement the above proof-of-concept simulations we here performed simulations varying parameter settings to address overall questions about variation, both between species but also within studies. We used the script from the delayed gratification task above and we performed four additional simulations. In case one we varied the level of exploration and ran simulations with *β*-values from 0.1 to 2 (cf. default value=1). In a second case we varied the learning rate of stimulus-response value learning using *α*-values from 0.2 to 2 (cf. default value=0.2). Thirdly we varied the size of the behavior repertoire to explore the effect of having a larger selection of behavior available. We varied the number of behaviors from 2 to 10. In the final case we used all default values (identical parameter values as in section Delayed gratification) to explore individual variation due the probabilistic nature of decision making. (Filename of script: 2b-Delayed-gratification-Case-study.txt. See notes in script on how to vary parameters.)

## Results

### Simulations of tasks from Animal AI Olympics

Overall, simulation results show that when subjecting a general associative learning mechanism, such as A-learning ([30, 34, 35]), to this array of tests it successfully learns most tasks. However, the nature of one task prevents this learning mechanism from extracting information from the current stimulus situation required for learning to take place (object permanence task). When adding a biologically plausible representation of an animal short-term memory to the learning algorithm that task was also solved successfully. See table 1 for key aspects needed to learn each respective task.

#### Two-choice mazes

In both T- and Y-mazes few behaviors need to be explored to find the food reward, and learning to choose the right arm develops quickly. After 10 trials the correct arm is chosen predominantly (Fig. 1A) and the probability to choose the correct arm increases from the first correct choice (Fig. 1B).

**Fig 1.**
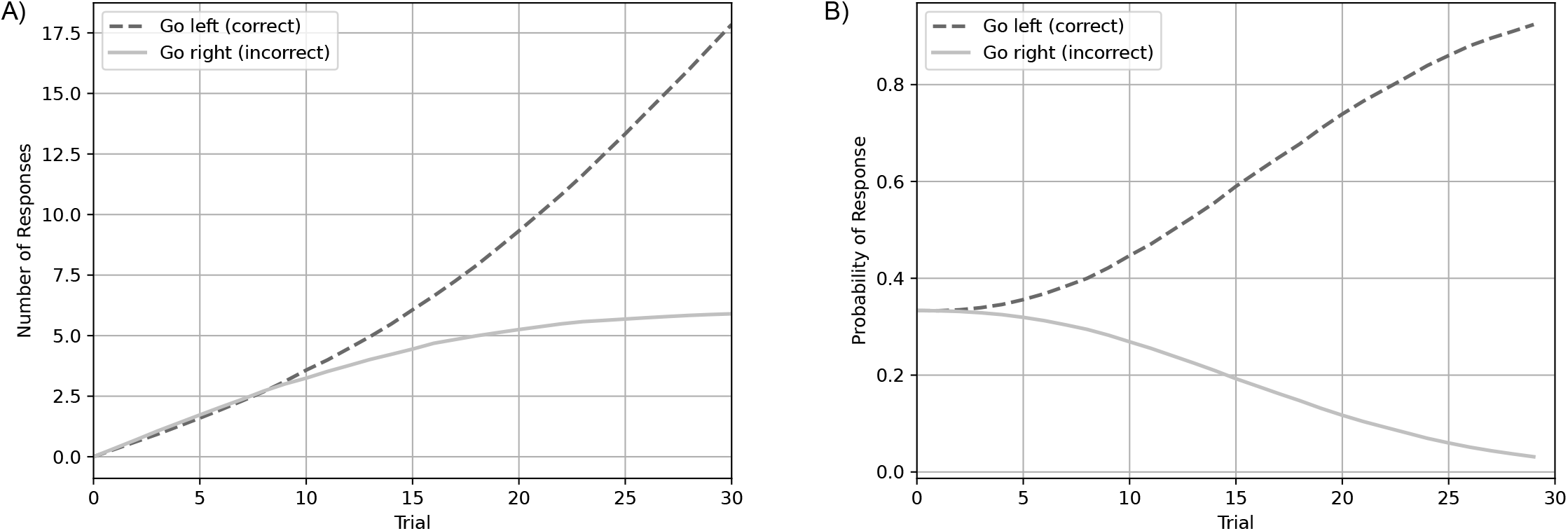
Results from two-choice mazes. A: Number of responses (cumulative) per trial, and B: probability of response per trial.

#### Delayed Gratification

The delayed gratification task requires inhibiting behaviors initially directed towards food, and this task cannot be learned until such an inhibiting behavior is first selected and has a positive outcome. This is shown by initial incorrect behavior both in terms of the number of incorrect behaviors (Fig. 2A) and that the likelihood of incorrect behavior at first grows rapidly (Fig. 2B). However, if the behavior repertoire of an organism includes some inhibiting behavior, such as waiting, then an associative learning mechanism can learn to delay a gratification if waiting is selected and results in some reward (Fig. 2). This is driven by the fact that stimulus-response value for waiting becomes larger than the stimulus-response value for taking the food immediately (see supplementary information for stimulus-response value figures). This way, self-control defined as suppressing an immediate drive in favour of delayed rewards, can emerge through associative learning [26, 51].

**Fig 2.**
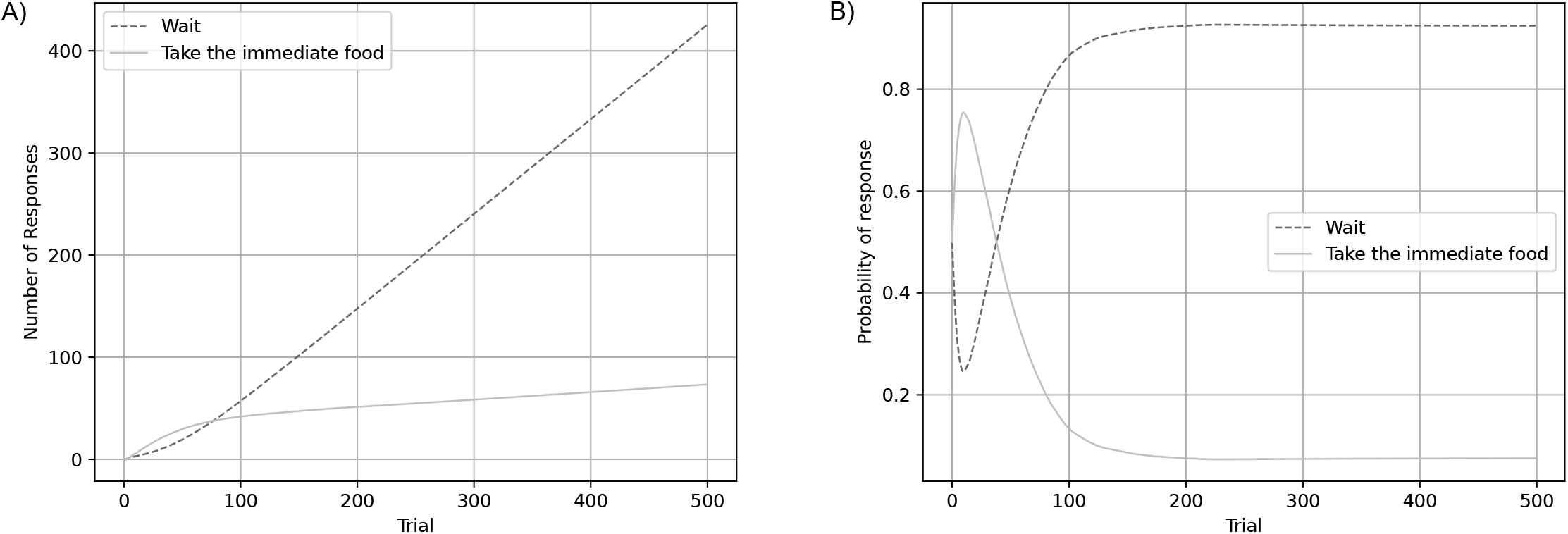
Results from delayed gratification. A: Number of responses (cumulative) per trial, and B: probability of response per trial.

#### Detour task

Like in delayed gratification tasks, for detour tasks to be solved some apparently non-rewarding behavior must first be explored, such as ignoring the sight of food or navigate around some barrier, and at a later time step produce some reward. Initially, incorrect behaviors are more common leading to few rewards in initial trials (Fig 3A) and initially higher probabilities for errors than correct behaviors (Fig 3B). However, results show than an associative learning mechanism can support solving detour tasks as long as detour behaviors are explored and have positive outcomes.

**Fig 3.**
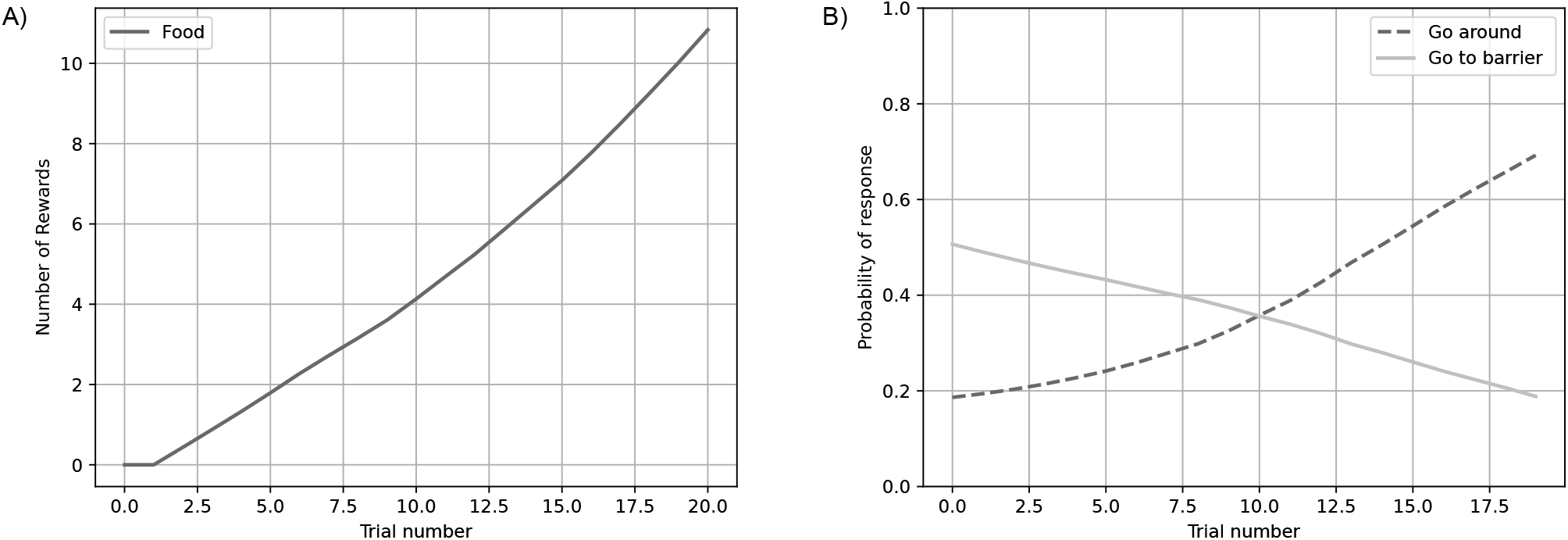
Results from detour task. A: Number of responses (cumulative) per trial, and B: probability of response per trial.

#### Cylinder task

Solving a cylinder task is related to both delayed gratification- and detour tasks in that reaching for inaccessible food must be inhibited to solve the task. However, a cylinder task is different in that it depends on initial training of a reward collecting behavior, that needs to be exhibited in the test phase. Initial training is transparent in that a correct behavior immediately results in a reward (Fig 4). Correct behavior in the test phase will increase in probability (Fig 4B) as soon as a correct behavior is selected, instead of the apparent incorrect behavior, reaching for the inaccessible food (see also [51] for training effects of self-control).

**Fig 4.**
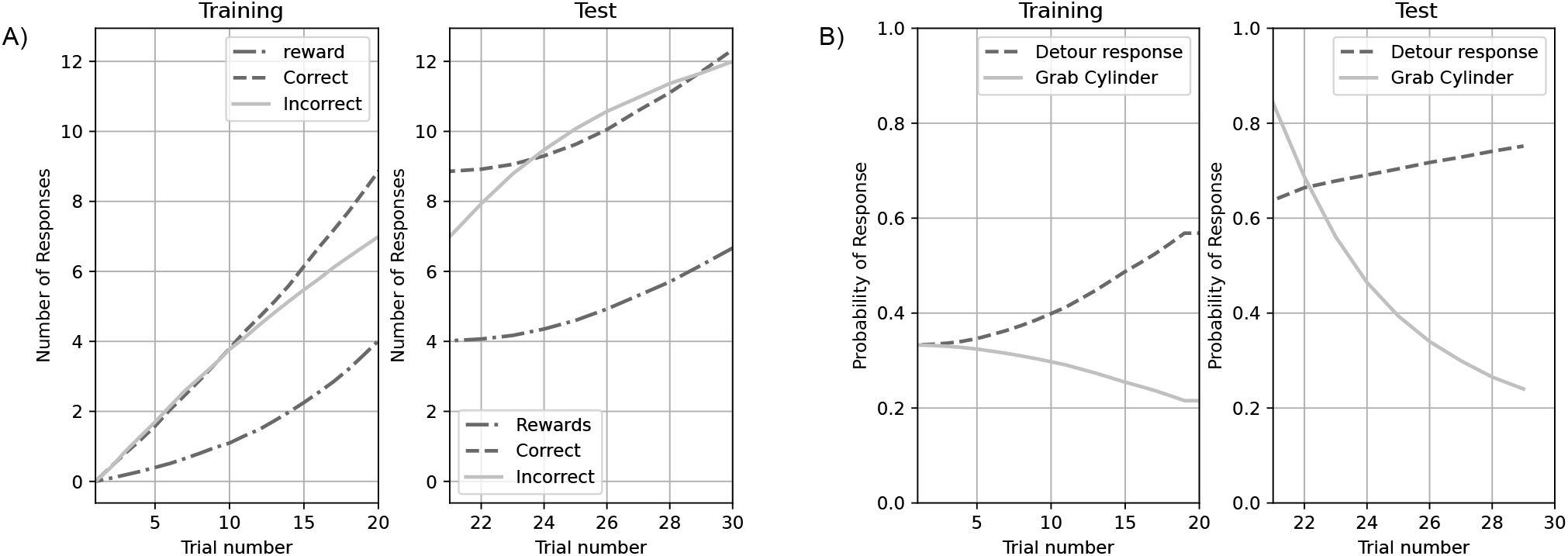
Results from cylinder task. A: Number of responses (cumulative) per trial, and B: probability of response per trial.

#### Thorndike’s Puzzlebox

To solve Thorndike’s box ‘k’ requires performing three behaviors, in no particular order, before a reward can be gained. The fact that correct behaviors are not immediately rewarding makes this task relatively difficult which is shown by the high number of behaviors that are required initially to solve this task (Fig 5A), and that the probability of performing the correct behaviors stabilize at relatively low levels (Fig 5B).

**Fig 5.**
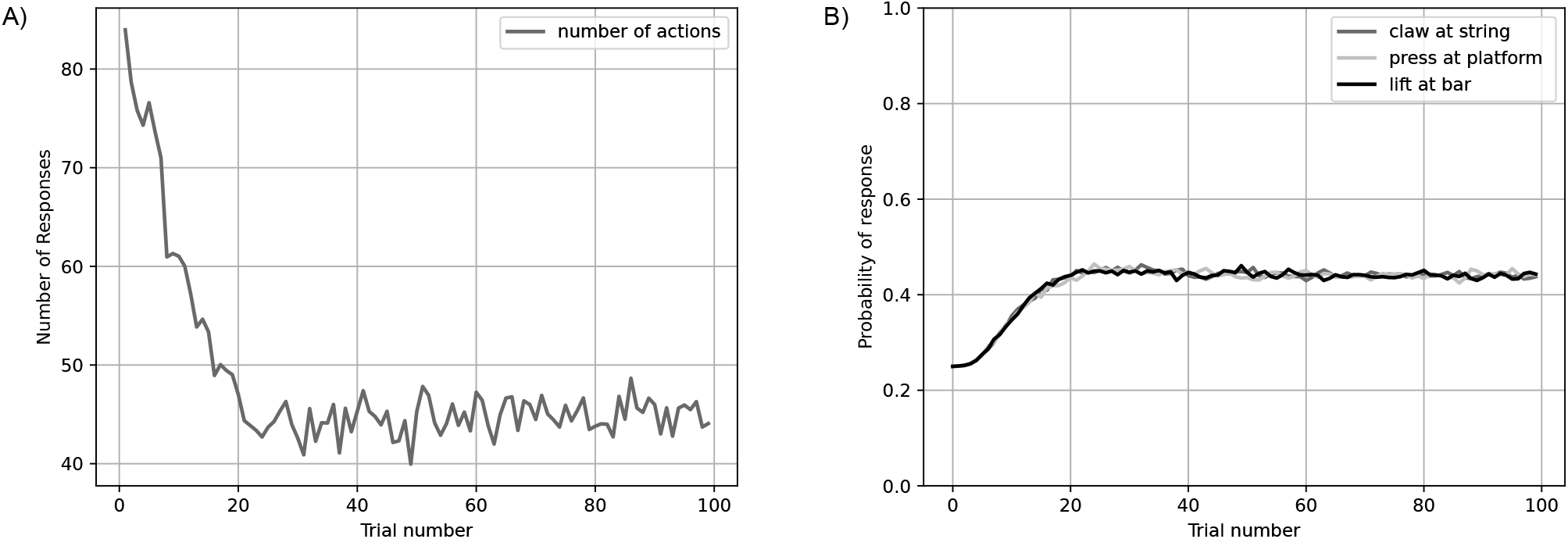
Results from Thorndike’s puzzlebox ‘k’. A: Number of responses (non-cumulative) per trial, and B: probability of response per trial.

#### Spatial Elimination

This task required choosing between two different stimuli, and a reward was presented immediately after a correct choice. This resembles the logical structure of the Y- and T-mazes, and the results are similar to that task, in that after approximately 10 trials the correct stimulus is predominantly chosen (Fig 6A) and the probability to choose the correct stimulus increases from the first correct choice (Fig 6B).

**Fig 6.**
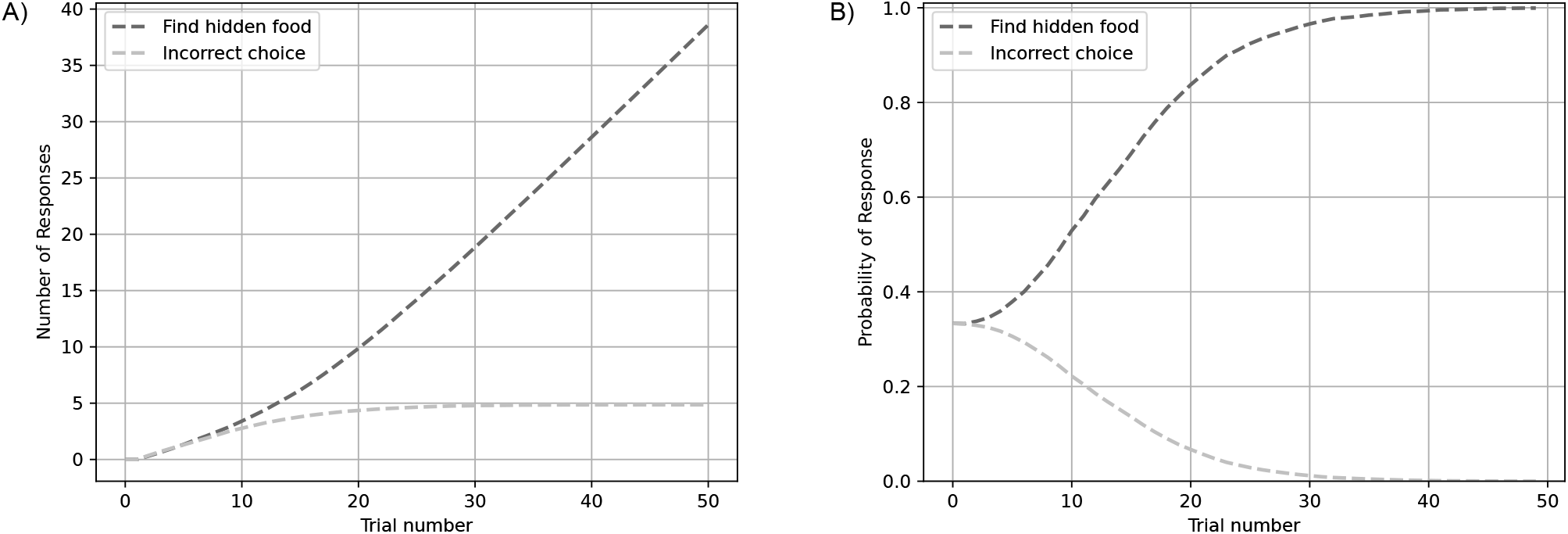
Results from spatial elimination. A: Number of responses (cumulative) per trial, and B: probability of response per trial.

#### Gravity Bias

The gravity bias task was simulated in three phases, first to see if it could be learned to collect a reward straight below where it was dropped, in line with what is expected from gravity (Fig 7A, D). Then a training phase was included for learning to collect food hidden behind doors (Fig. 7B, E). Finally, a test phase was included to see if a new behavior, that did not adhere to gravity as in phase 1, could be learned (Fig 7A, D). Simulations show that a gravity bias can emerge from associative learning (Fig 7D) and that a test in conflict with a previously learned gravity bias can also be mastered (Fig 7F). In phase 2, the new behavior to get food in boxes was crucial for the test in phase 3. However, note that training in phase 2 has no effect on the previously learned gravity bias, that is later altered in phase 3.

**Fig 7.**
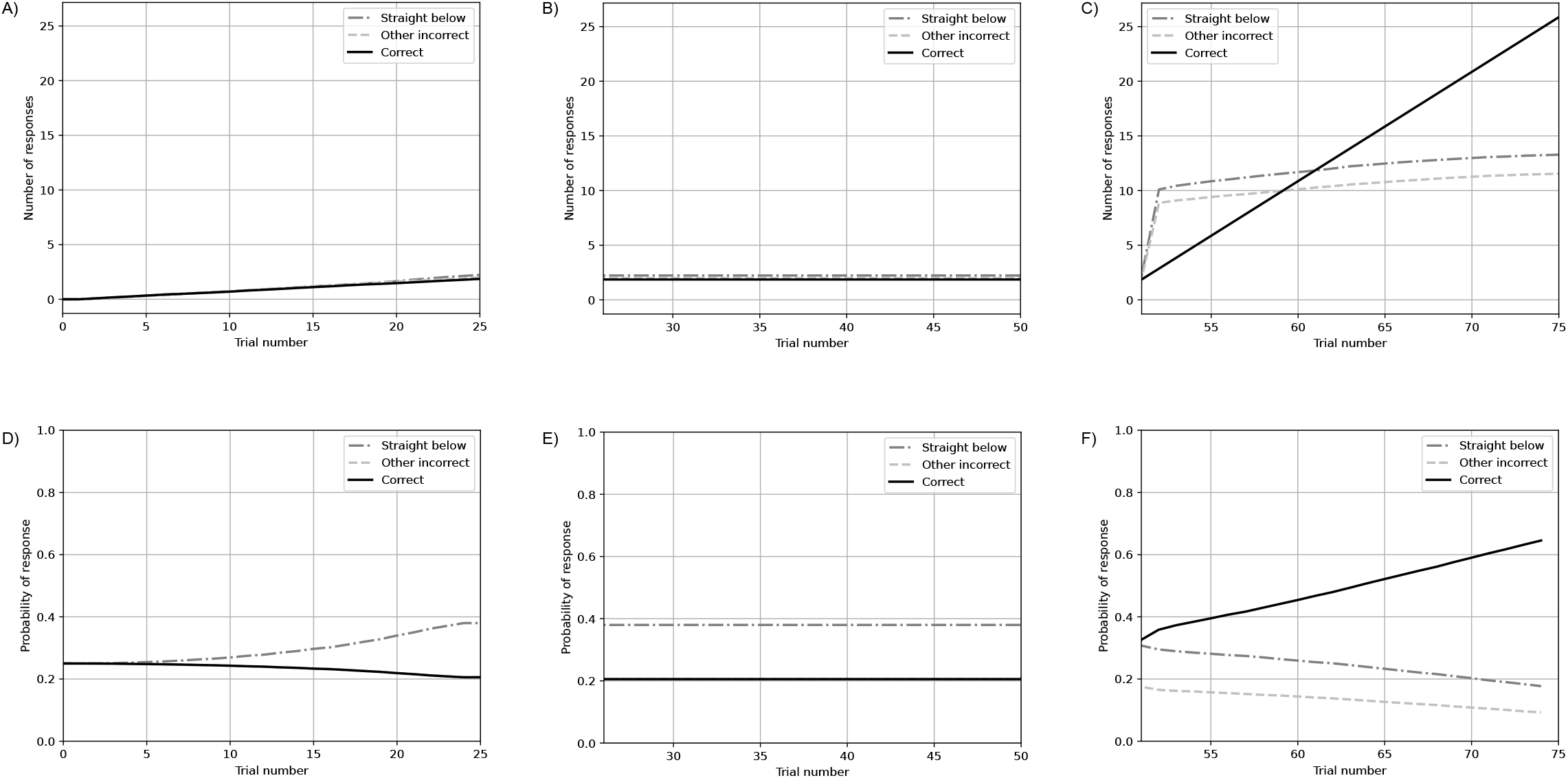
Results from gravity bias. Number of responses (cumulative) per trial for simulation of A: phase 1, B: phase 2, and C: phase 3. D: Probability of response per trial for the simulation of D: phase 1, E: phase 2, and F: phase 3.

#### Radial Maze

Although the choice situation is identical when returning from each of the eight arms, the task can still be learned in several ways. Overall, choosing to move an uneven amount of steps to the left or right (−3, −1, +1,or +3) was learned, and this is the quickest way to collect all food hidden in the maze (Fig 8B). This solution to the problem has been called “algorithmic” behavior [49] and these simulations show how such learned patterns of movement emerge through associative learning. That learning takes place is also shown by the reduction in number of arm visits per trial (Fig 8C). It may seem counter-intuitive that even steps (−2 and +2) are the most frequent responses (Fig 8A). However, this comes from even steps being frequently rewarding but at the same time preventing subjects from finishing a trial. In addition, as the maze had eight arms with one reward in each arm, going straight and returning back to the arms visited last were the least chosen behaviors. To show more clearly that “algorithmic behavior” develops through associative learning we included a figure with an individual run(Fig 8D). This shows how one behavior quickly can become a dominant strategy (Fig 8D).

**Fig 8.**
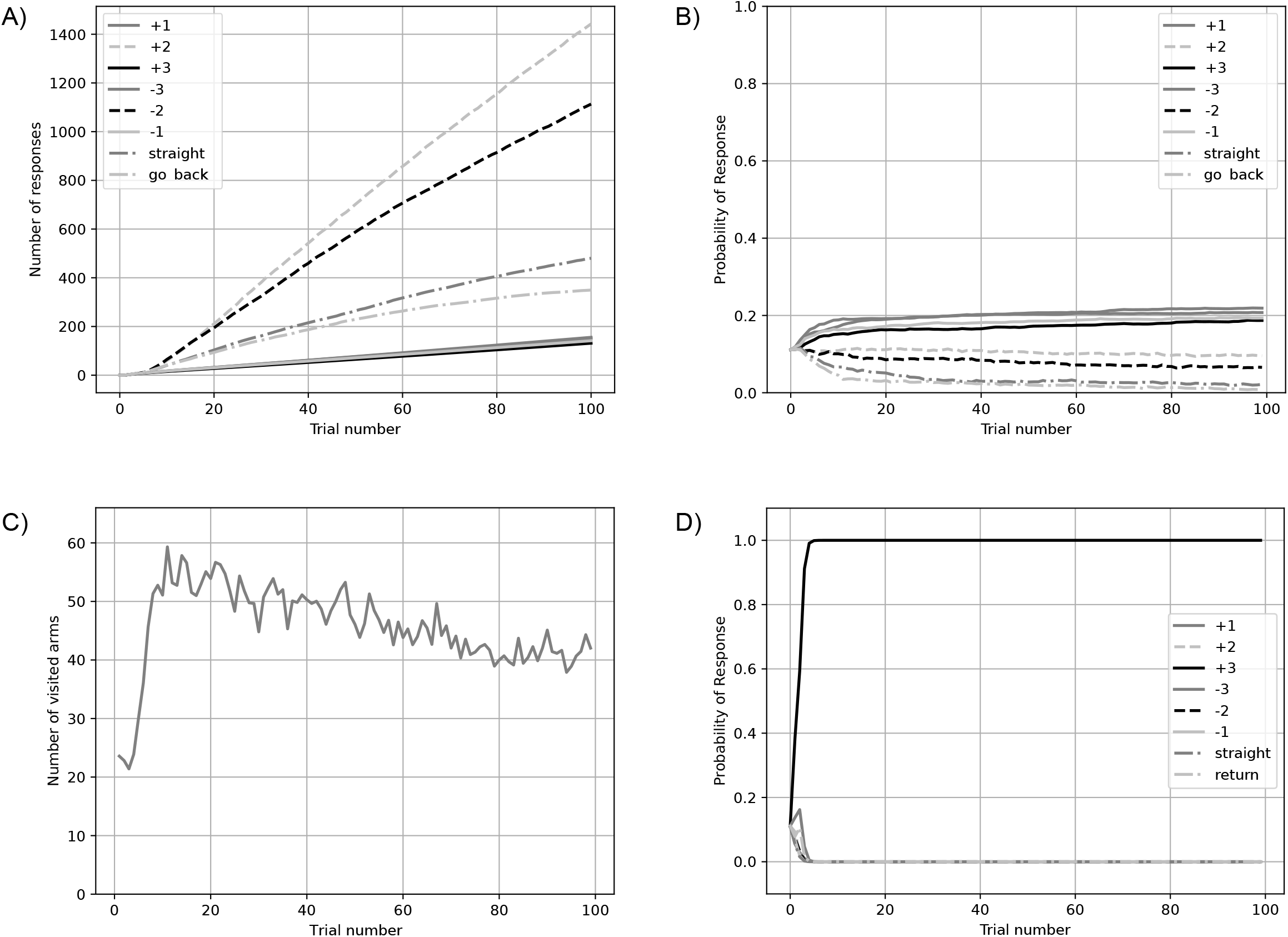
Results from the radial maze simulation. A: Number of responses (cumulative) per trial for simulation of 500 subjects. B: Probability of response per trial for the simulation of 500 subjects. C: Number of visited arms per trial for 500 subjects. D: Probability of response for a single subject.

#### Object Permanence

This task involved a sequence of steps prior to the choice situation where three large cups were present. This task could not be solved above chance level by the associative learning mechanism alone because the choice situation was identical irrespective of where the reward was located. The only discriminative information available was in a time step prior to the choice situation. However, when using an integrated memory (a standard animal short-term memory ([11, 37, 38, 40]) that provides the associative learning mechanism with information about the time step prior to the choice situation, the task could be solved. Correct behaviors became more common than errors after some 100 trials (Fig 9A) and the probability of choosing the correct cup increased well above chance levels before 100 trials (Fig 9B).

**Fig 9.**
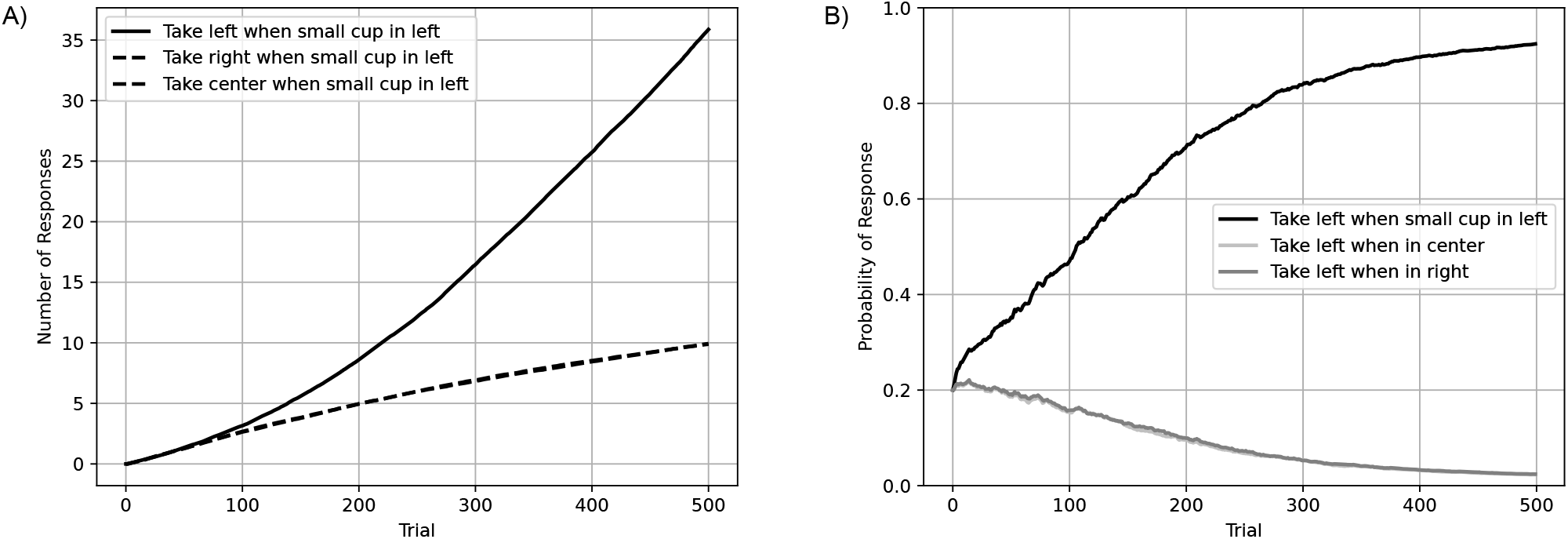
Results from object permanence. A: Number of responses (cumulative) per trial, and B: probability of response per trial for the object permanence simulation

#### Numerosity

In this numerosity task, a two choice situation between bowls with different quantities of food, from one to six, was presented. The associative learning mechanism developed a preference rapidly for the bowl with the largest number of food items (Fig. 10A, B). In general, associative learning is expected to result in preferences for the largest quantities of food as long different quantities can be told apart, and A-learning will given sufficient information result in optimal behavior ([34]).

**Fig 10.**
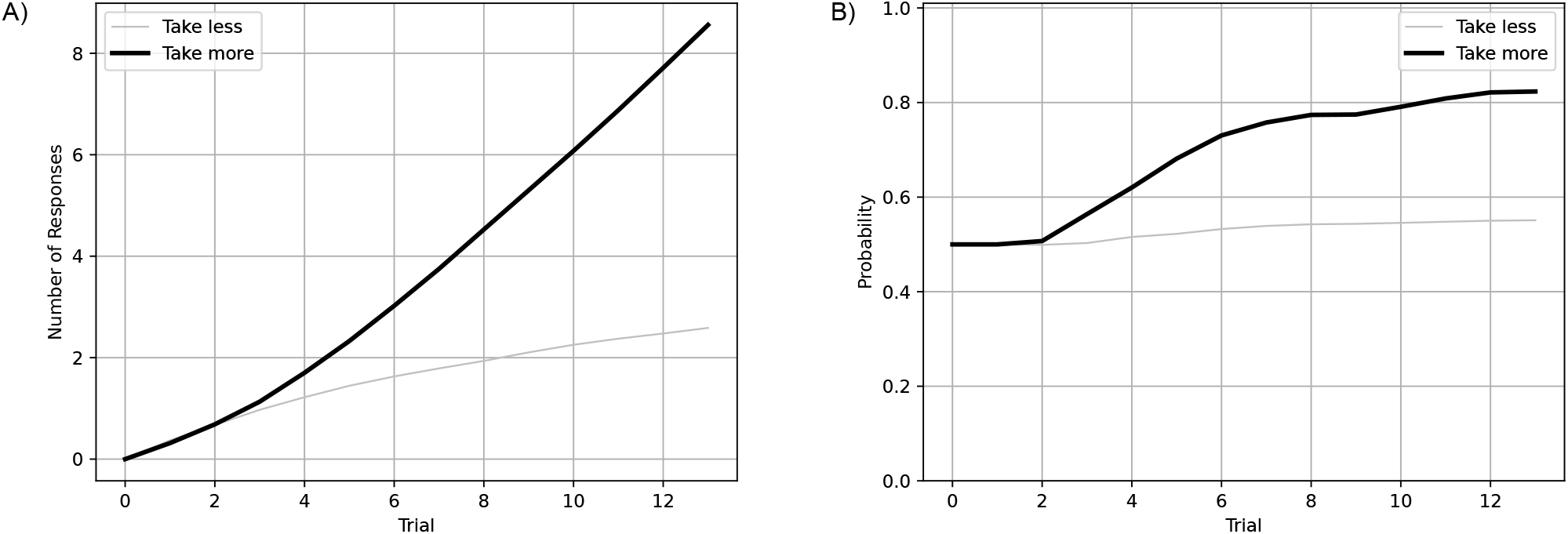
Results from numerosity. A: Number of responses (cumulative) per trial, and B: probability of response per trial.

#### Tool use

In this tool task the problem was to choose between two cloths that could be pulled to get a reward. One cloth had food on top of it whereas the other cloth had food next to it. Results show that learning to choose the cloth with food emerge after some 20 trials (Fig 11A), and that the probability of ignoring the cloth without food increased rapidly as it did not yield a positive outcome (Fig 11B).

**Fig 11.**
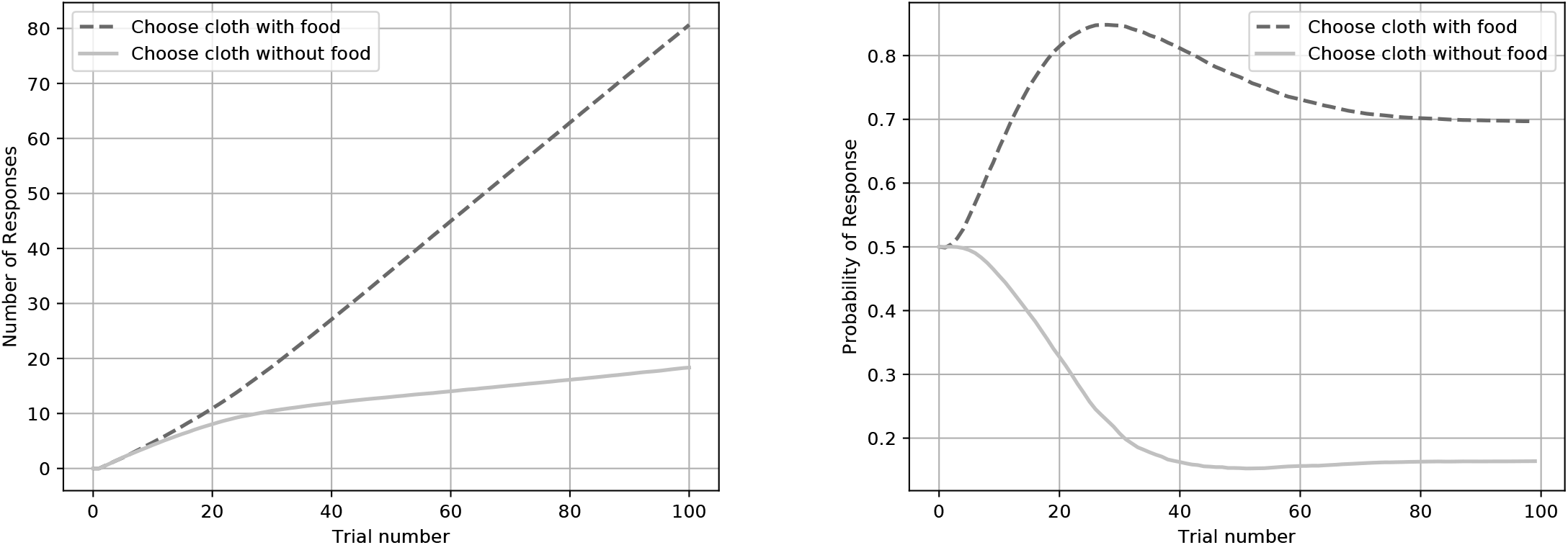
Results from tool use. A: Number of responses (cumulative) per trial for the tool use simulation, and B: probability of response per trial.

### A case study on variation in learning and test performance

All previous simulations were performed using identical parameter values. Here we show the impact of varying parameter values on learning, using the script from the Delayed gratification task above. First we varied the f3-parameter that regulates exploration. A low value makes it less likely that the behavior with the highest estimated value is chosen, allowing previously non-rewarding, or only little rewarded, behavior to be exhibited^1^. Results show that learning to delay the gratification, that is exert some self-control and learn to wait in the presence of a reward, can be learned at intermediate and high levels of exploration (low *β*-values). But, lower levels of exploration (higher *β*-values) results in choosing immediate rewards because they were initially rewarding and low levels of exploration makes it less likely to choose non-rewarding behaviors. Too little exploration prevents learning to wait for a larger reward as the waiting behavior is initially non-rewarding and becomes a hurdle that can only be overcome with higher levels of exploration (Fig 12A). We then varied the learning rate, the *α*-value that determines the update rate of stimulus-response values. Varying the *α*-value shows that learning can potentially be very rapid (Fig 12B). As a third step we varied the behavior repertoire size. This simply shows that as more behavioral options are available, all else being equal, more learning time is necessary for productive behavior to develop because as there are more behaviors to choose finding a rewarding behavior is less likely (Fig 12C). Even small increases in behavior repertoire size can prolong learning a behavior. Finally we performed the same simulation using only five subjects, showing that the probabilistic nature of decision making can have a large impact on performance with some subjects finding rewards rapidly making learning possible, whereas other individuals do not even learn after several hundreds of trials (Fig 12D). This mechanism can generate different behavior outcomes purely by chance. Please note that performance in all tasks above will vary similarly to how performance varied here.

**Fig 12.**
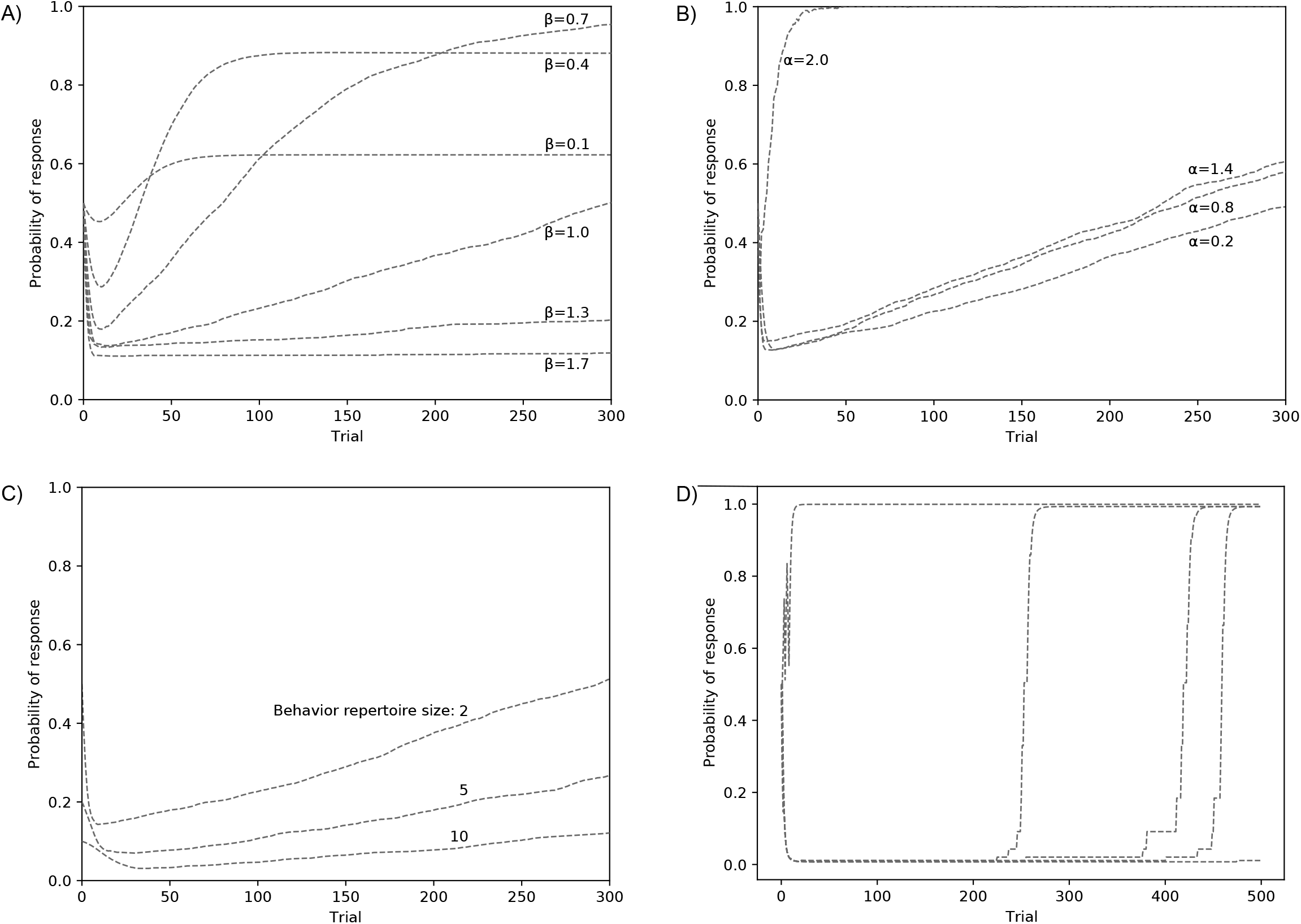
Results from the case study. These simulations were based on the Delayed gratification-task script. A: Probability of performing wait behavior as a response to food with different values for exploration. B: Probability of performing wait behavior as a response to food with different values for memory update rate of behaviors. C: Probability of performing wait behavior depending on behavior repertoire size. D: Variation between five individual runs with identical parameter settings.

Learning can be quicker, in some cases even instantaneous after one successful trial, and slower dependent on parameter settings in exploration, learning rate, and behavior repertoire.

## Discussion

Our computer simulations show that associative learning as a mechanism can potentially solve the animal intelligence tests selected by Crosby et al. [23, 24]. It is not surprising that relatively simple tests such as T- and Y-mazes are mastered. However, our simulations prove associative learning to be a powerful mechanism, that can solve problems of higher complexity. In general, as long as rewards can be found and tasks allow backwards chaining, associative learning can be sufficient for mastering tasks [34]. In backwards chaining, the final step yielding a reward is learned first. Subsequently, a neutral stimuli that precedes a food reward can acquire conditioned reinforcement value through stimulus value-learning and drive the learning of a behavior sequence [6, 34]. That associative learning can give rise to behavior sequences, makes apparently complex behaviors possible and enable animals to master tasks that are generally considered complex. For example the delayed gratification-, cylinder-, and tool use task. This corresponds well with analyses showing how tool-use [34], planning with self-control [26], and social learning [35] can emerge through associative learning. Although associative learning is often considered too simple for explaining animal intelligence in general [22, 52–55], modern models of associative learning are more powerful and more cognitive, than 50 years ago [56, 57]. Our results, together with current progress in both animal and AI learning [28–30, 58], makes it increasingly difficult to reject the idea that associative learning is a general mechanism that underlies animal intelligence.

The A-learning model alone could not solve the object permanence test because at the time of decision, no discriminatory information about the reward was available (see section on Object Permanence). However, animals remember arbitrary stimuli at least in the magnitude of seconds to a few minutes [40]. Thus, by adding a well described trace memory [37, 38] to the A-learning model, information becomes available at the final choice point, and the task can be solved. This shows that expanding associative learning models with other parts of behavior systems, such as a trace memory, is a tangible way for future research into animal intelligence. Internal variables, such as memory and motivational states, can potentially provide information to associative learning just like external variables.

The fact that some animals, obviously capable of associative learning, fail to learn some of these tasks may seem to contrast with our overall results. However, this formulation of associative learning generates considerable variability, which is potentially consistent with the variation that is observed between species (Fig 12). We addressed three factors that can cause variation in performance. First, in order to learn something new, previously non-rewarding behaviors must be tried out. Exploration varies greatly between species, and species less inclined to try new behaviors will in general suffer from lower performance [59, 60]. Low levels of exploration can even completely prevent the learning of a new task (Fig 12A). The costs of curiosity and high levels of exploration should, however, not be ignored. Exploring too much comes with costs, for example by wasting limited resources and through putting an individual in dangerous situations. Levels of exploration are likely a compromise between contrasting selection pressures that vary between species. Secondly, learning rates of both stimulus-response values (Fig 12B) and stimulus values will affect learning speed. Even rapid insight-like ‘one shot learning’ is possible through associative learning. Finally, all else equal, a greater behavior repertoire size makes it less likely to find a new solution to a problem (Fig 12C). A larger behavior repertoire comes with higher learning costs and requires certain circumstances to be favorable, for example a longer juvenile period as seen in great apes [61]. A-learning integrates genetic predispositions with general learning processes and can, therefore, account for between species variation and questions of domain-specific adaptations [34]. However, detailed explanations of empirical results are beyond the scope of this study.

Due to the stochastic nature of decision making, individual simulation runs can differ greatly in performance (Fig 12D). This is in line with findings that choice behavior is affected by stochastic processes [62, 63] and may provide a null hypothesis for tests of animal intelligence. This provides a reasonable alternative to interpretations of individual variation as caused by individual differences in intelligence or unidentified contextual variables [64]. The fact that some individuals fail whereas others do not may not be caused by some individuals being smarter than others, as it is often suggested [65–68]. However, as mentioned above, the level of exploration may vary consistently between individuals and affect performance in problem solving [69].

How do these results inform the general process vs. adaptive specialization debate? Like others [4, 26, 30, 34, 35, 56, 57], we conclude that associative learning is an underestimated general mechanism that can account for a large array of behavioral phenomena observed in animals. Take the numerosity task, here associative learning may provide an alternative to the hypothesized specialized module for numerosity, the approximate number sense [70, 71]. However, this suggestion requires further analyses (see e.g. special issue on the origin of a number sense [5]). Furthermore, our results favor the idea that species differences emerge from genetic differences in general processes rather than due to adaptive specialization of separate cognitive traits each with its own unique evolutionary trajectory. Assumptions of adaptive specialization may inflate estimates of cognitive traits subject to convergent evolution. For instance, it is not self evident that similarities in how ravens and great apes solve similar tasks means that they independently evolved similar cognitive mechanisms, when an associative learning account is plausible [26, 72, 73]. We do not, however, reject the idea of adaptive specialization of cognition. If the development of some behavior does not adhere to descriptions of associative learning, they may be better described in terms of adaptive specialization. This may include phenomena where a high performance is required from scratch, for example navigation in desert ants [74], and when first-time migratory birds use celestial cues for flight direction [75] or the Earth’s magnetic field for foraging decision [76].

## Conclusion

In conclusion; we can not reject that current models of animal learning may suffice to generate flexible and intelligent behavior in animals, with learning being guided by species-specific behavior systems. Interestingly, models in animal psychology and machine learning share many similarities [30]. As long as motor/perceptive systems are accounted for (as in our simulations), associative learning mechanisms can solve apparently complex tasks. This aligns well with results from the Animal-AI Olympics [24]. Agents generally did not struggle with learning about the value of rewards and how to interact with the environment. Instead they were limited by not having the appropriate perceptive/motor systems. For example, many agents struggled to do well on the tool-use tasks, because they were not equipped with the right code to be able to manipulate objects. So, perhaps it is not the sophistication of learning algorithms per se that limits intelligent behavior of AI systems. Take, for example, a 10 gram bird that has evolved the ability to use starry nights for migration, or a bolas spider that makes bolas at the end of a silk thread and uses this as a lasso to catch prey. Here, evolution has resulted in integrated learning-, motor-, attentional- and perceptual capacities that together make up autonomous organisms that we still struggle to describe. Therefore, the challenge for AI to approach animal-like intelligence may not lie in creating new, clever algorithms, but in understanding the complexity of biological systems in terms of perception, motor flexibility, and behavior regulation. To further progress, future research could benefit from integrating embodied aspects of cognition [77, 78] with a general mechanism of associative learning.

## Supporting information

**S1 Appendix** Includes information on the simulator scripts and where to find them, as well as additional figures belonging to the simulations discussed in the paper.

## Acknowledgments

We would like to thank Markus Jonsson and Magnus Enquist. JL was supported by Knut and Alice Wallenberg Foundation (KAW 2015.005).

## Supplementary information for

### 1 Scripts and Learning Simulator

All scripts needed to run the computer simulations are available as individual text files and can be downloaded at https://doi.org/10.17045/sthlmuni.17068409. The scripts can be run in the Learning Simulator [1] available at https://www.learningsimulator.org. For details see documentation at https://learningsimulator.readthedocs.io.

### 2 Additional plots of all Simulations

To complement figures in the main manuscript we here present additional figures of all learning tasks. These additional figures show changes in stimulus-response values *v*(*s* → *b*) in the different tasks.

#### 2.1 Two-choice Mazes

**Figure 1:**
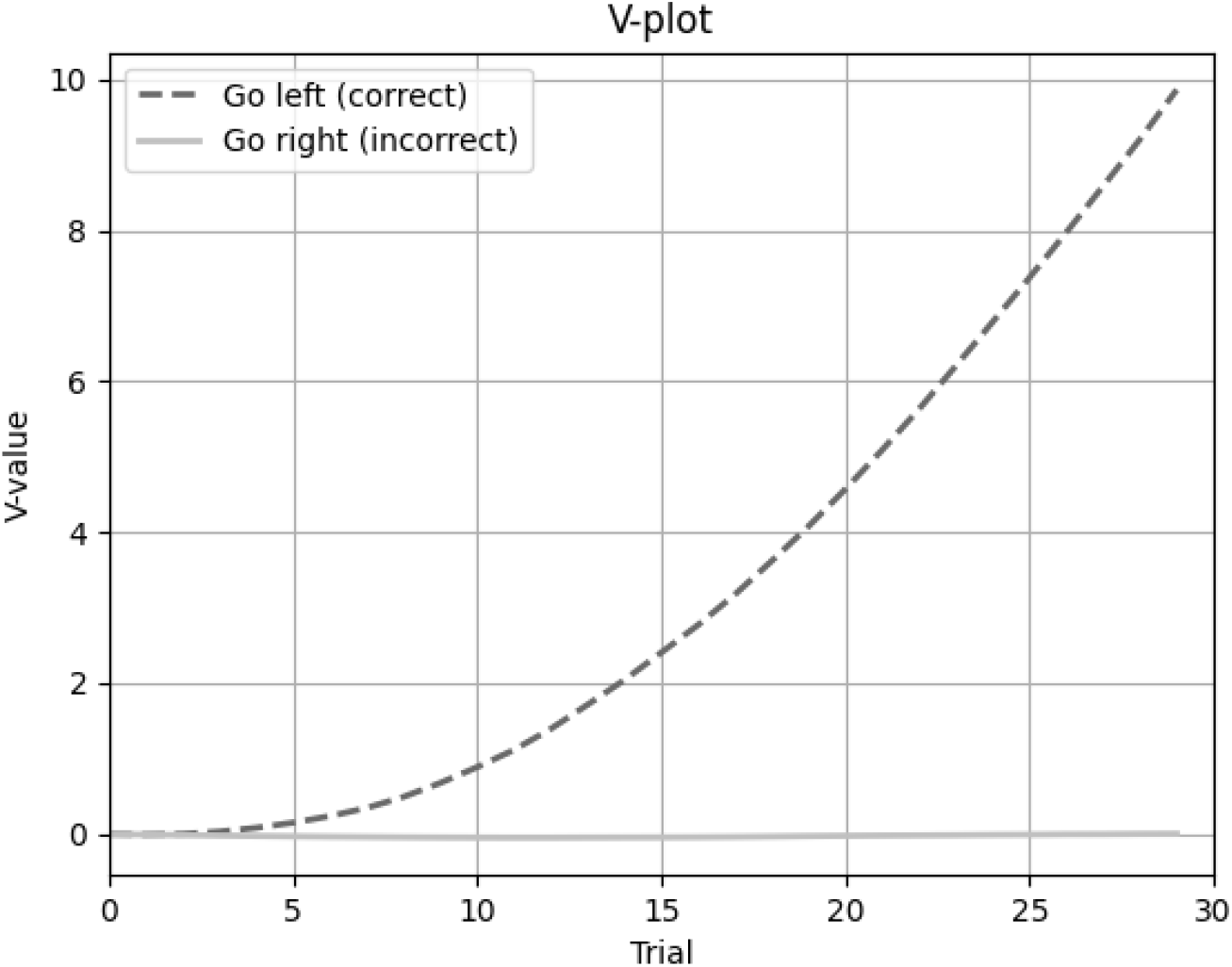
V-plot for the two-choice mazes simulation for two behaviors at *s*(start)

#### 2.2 Delayed Gratification

**Figure 2:**
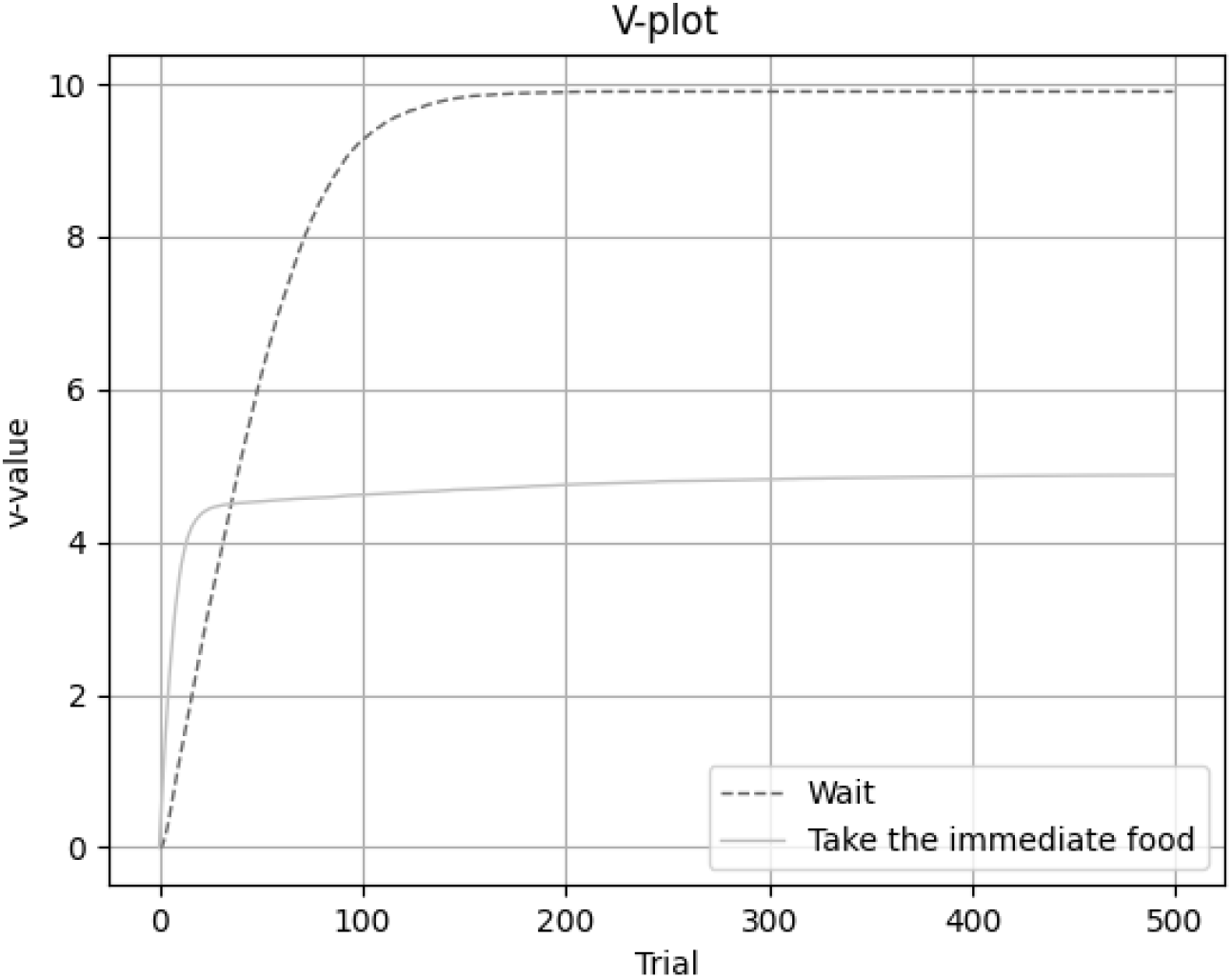
V-plot for the delayed gratification simulation for two behaviors at *s*(immediate food)

#### 2.3 Detour Tasks

**Figure 3:**
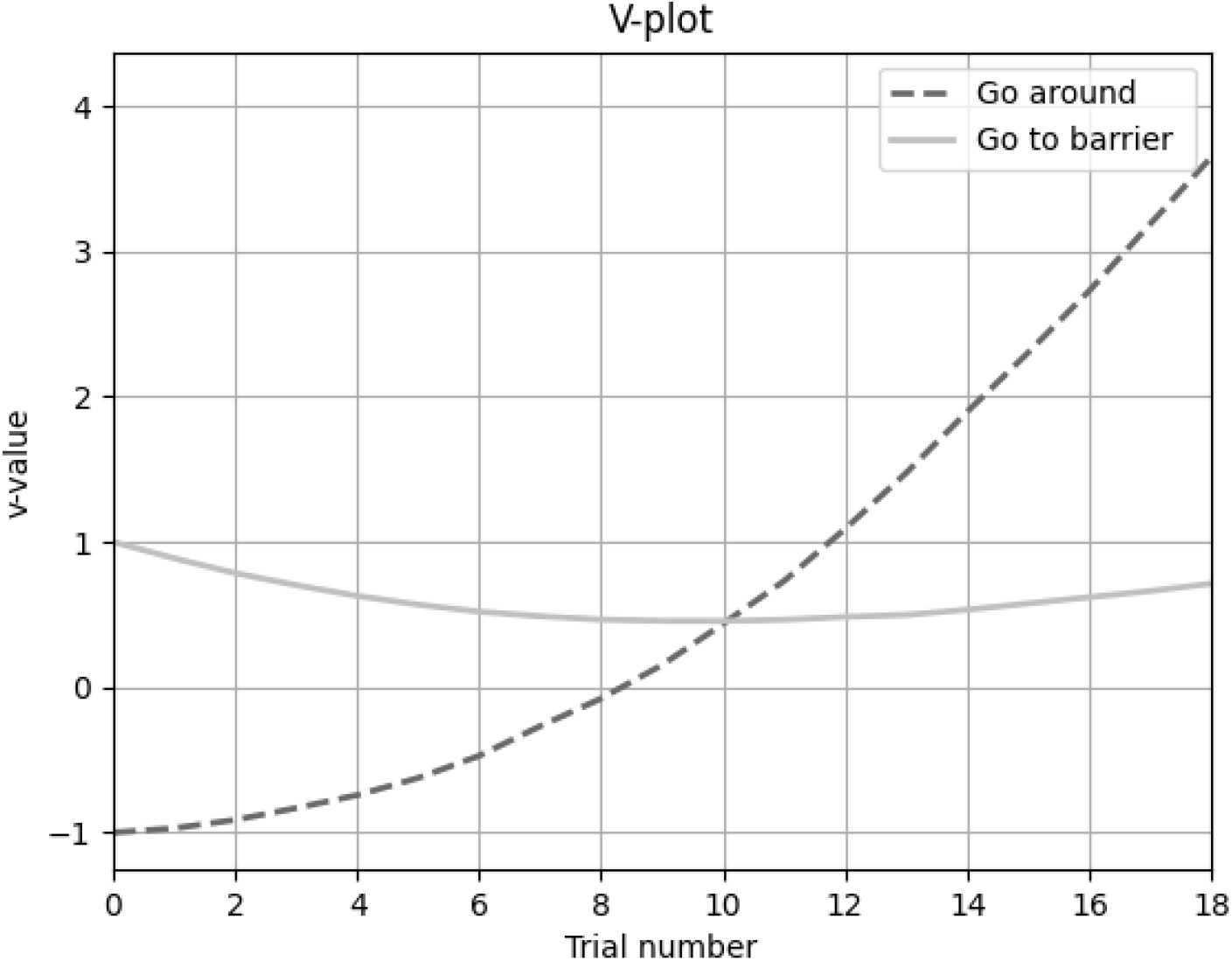
V-plot for the detour task simulation for two behaviors at *s*(start).

#### 2.4 Cylinder Tasks

**Figure 4:**
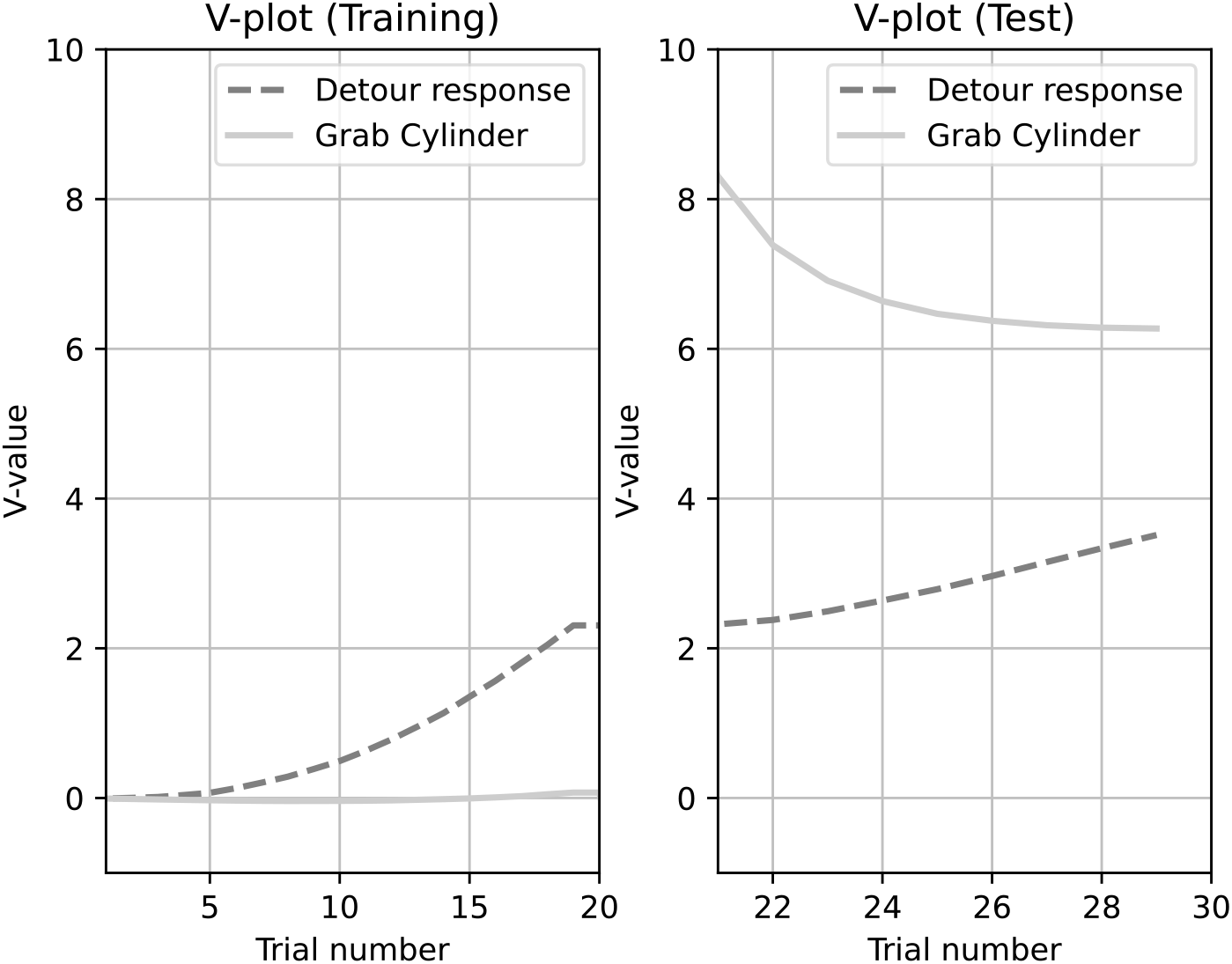
V-plot for the cylinder task simulation for *s*(cylinder) → *b*(detour response) and *s*(food inside) → *b*(grab cylinder)

#### 2.5 Thorndikes Puzzlebox

**Figure 5:**
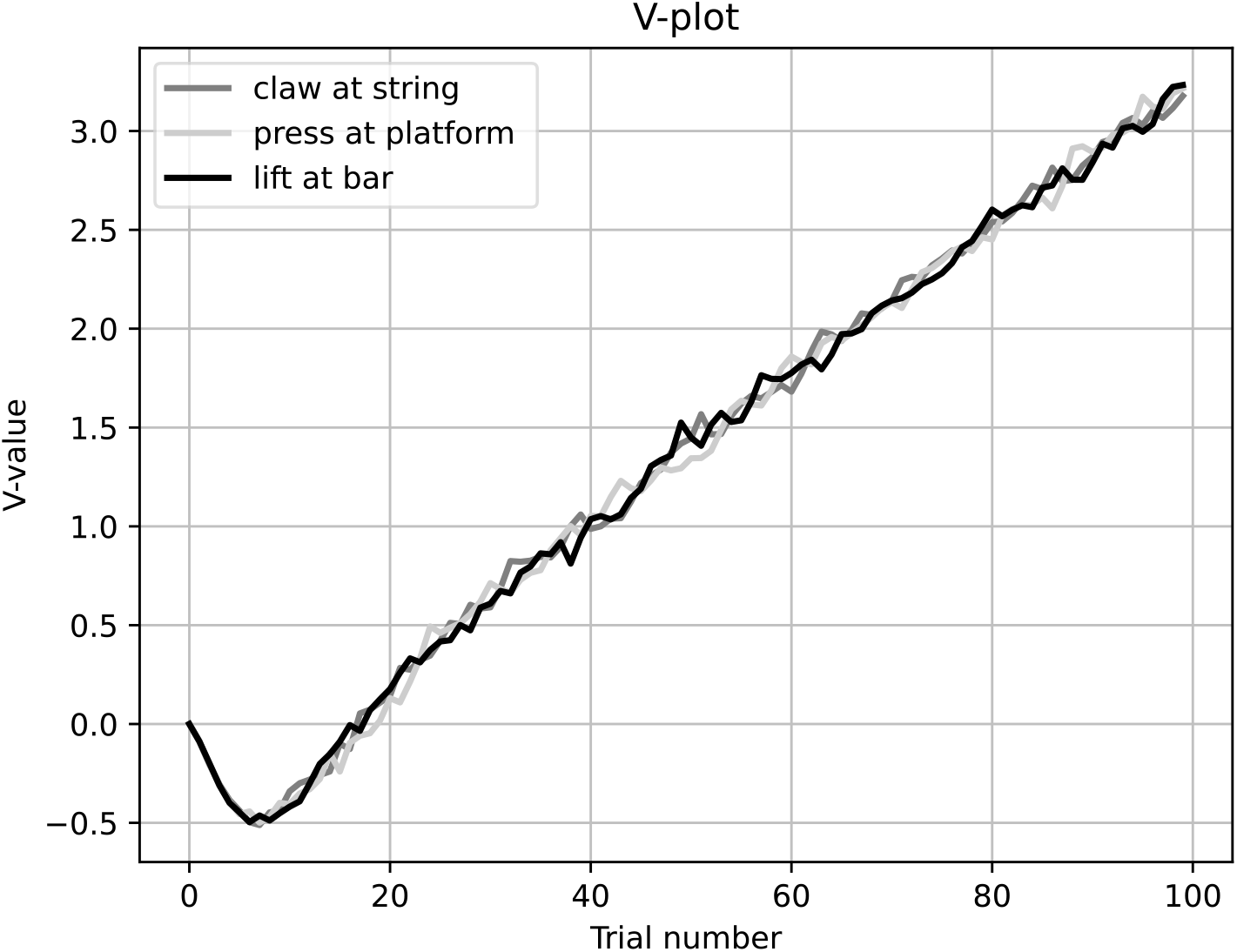
V-plot for the Thorndikes puzzlebox simulation for *s*(string) → *b*(claw) *s*(platform) → *b*(press), and *s*(bar) → *b*(lift).

#### 2.6 Spatial Elimination

**Figure 6:**
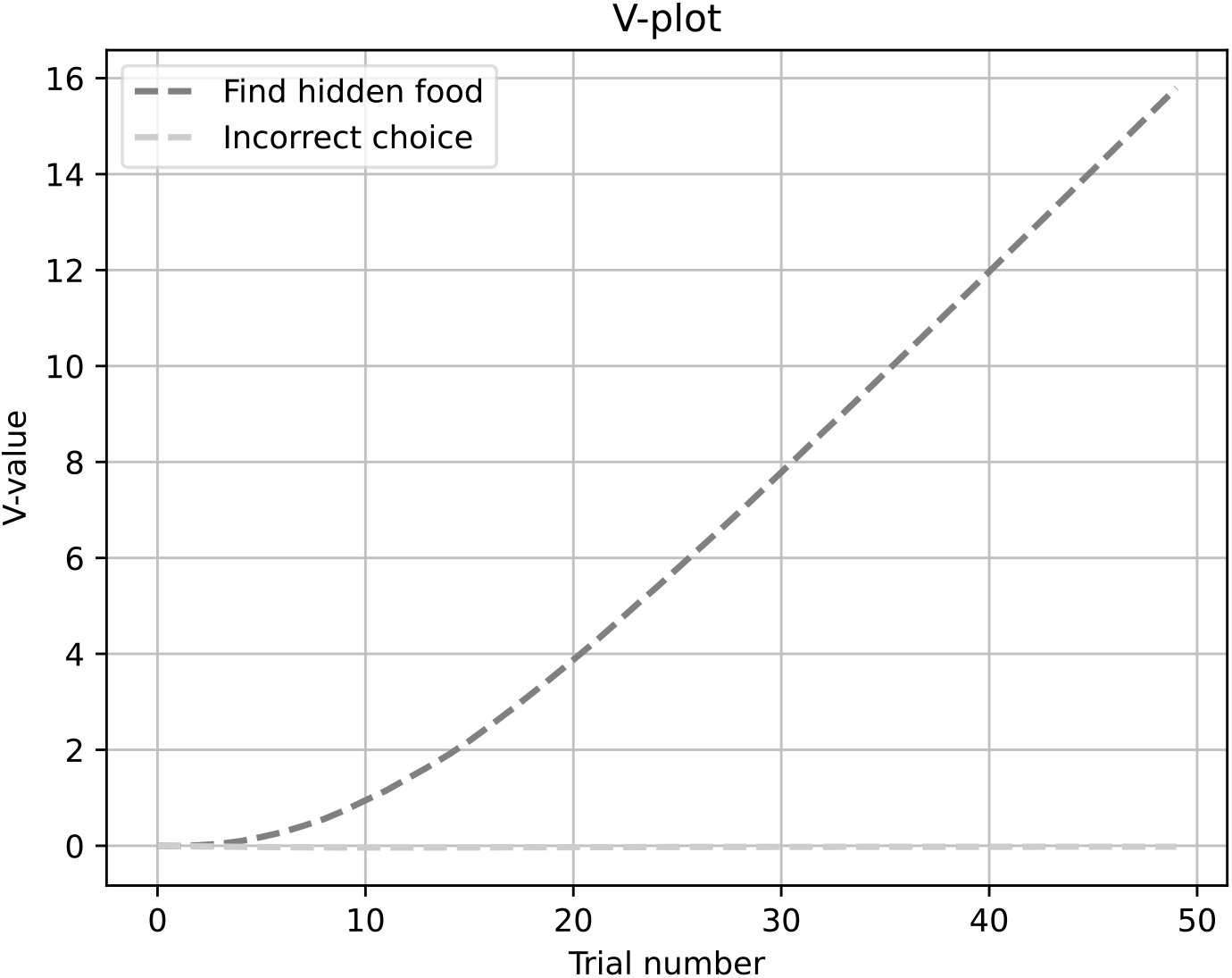
V-plot for the spatial elimination simulation for *s*(inclined board) → *b*(choose inclined) and *s*(flat board) → *b*(choose flat).

#### 2.7 Gravity Bias

**Figure 7:**
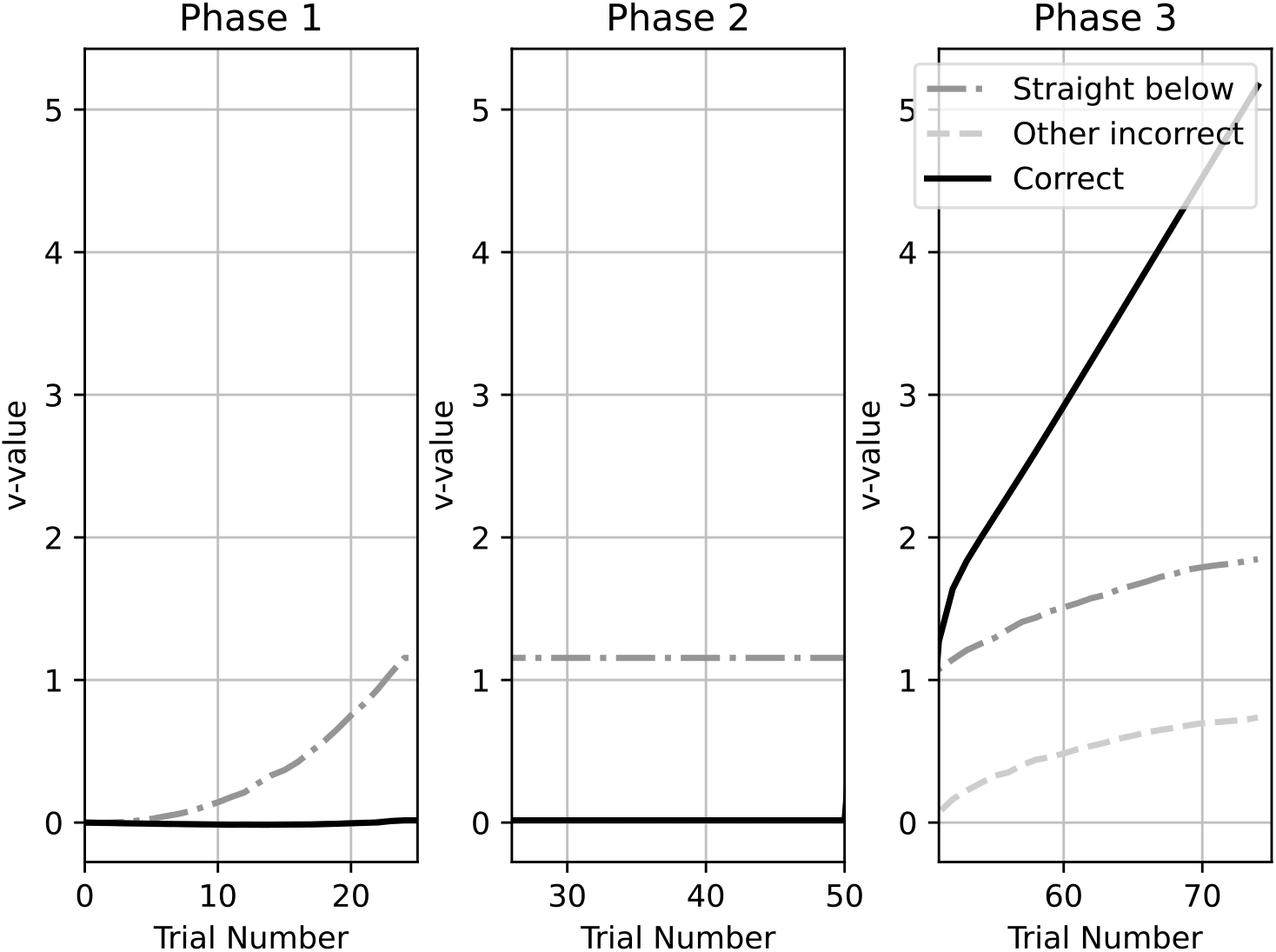
V-plot for the 3 phases of the gravity bias simulation.

#### 2.8 Radial Maze

**Figure 8:**
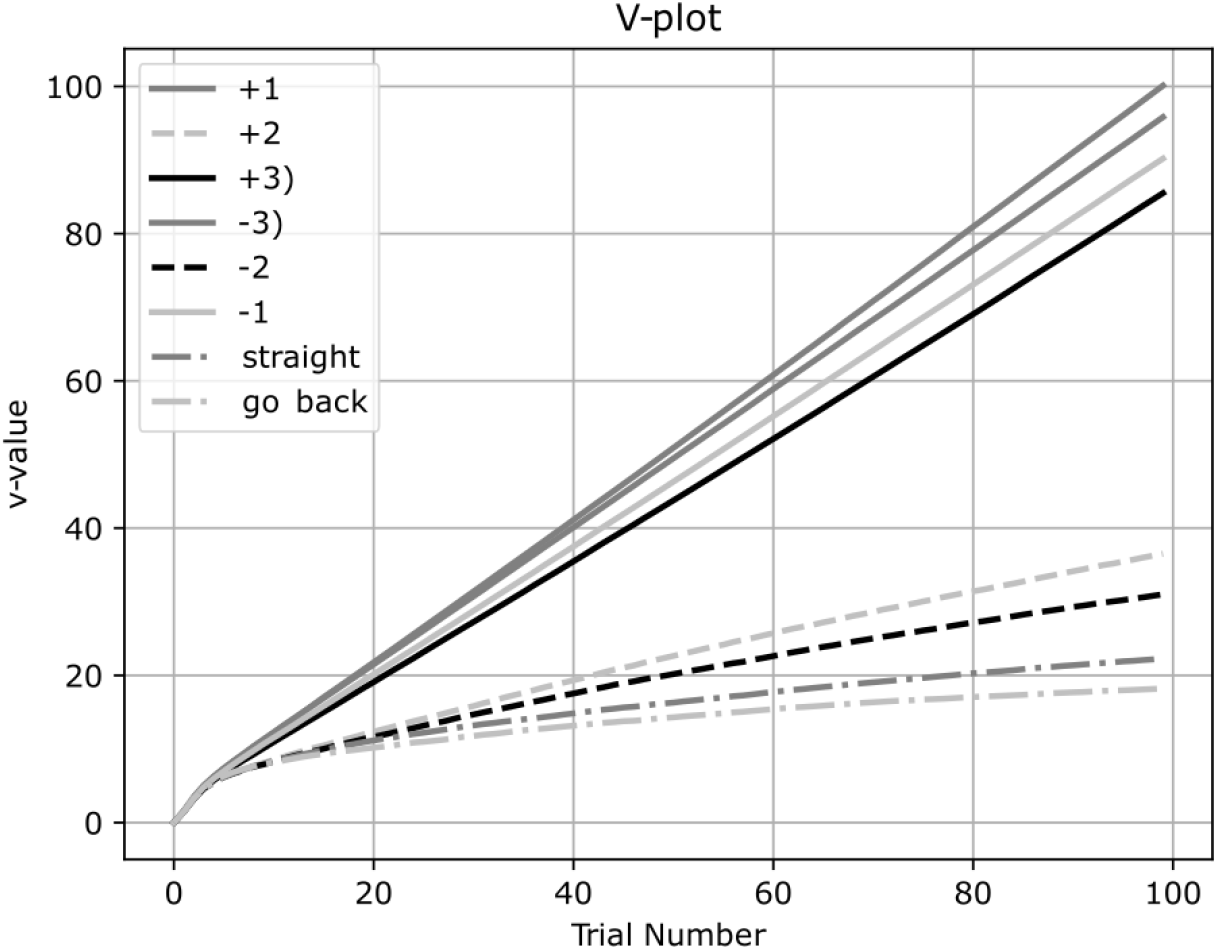
V-plot for the radial maze simulation of behaviors at *s*(center).

#### 2.9 Object Permanence

**Figure 9:**
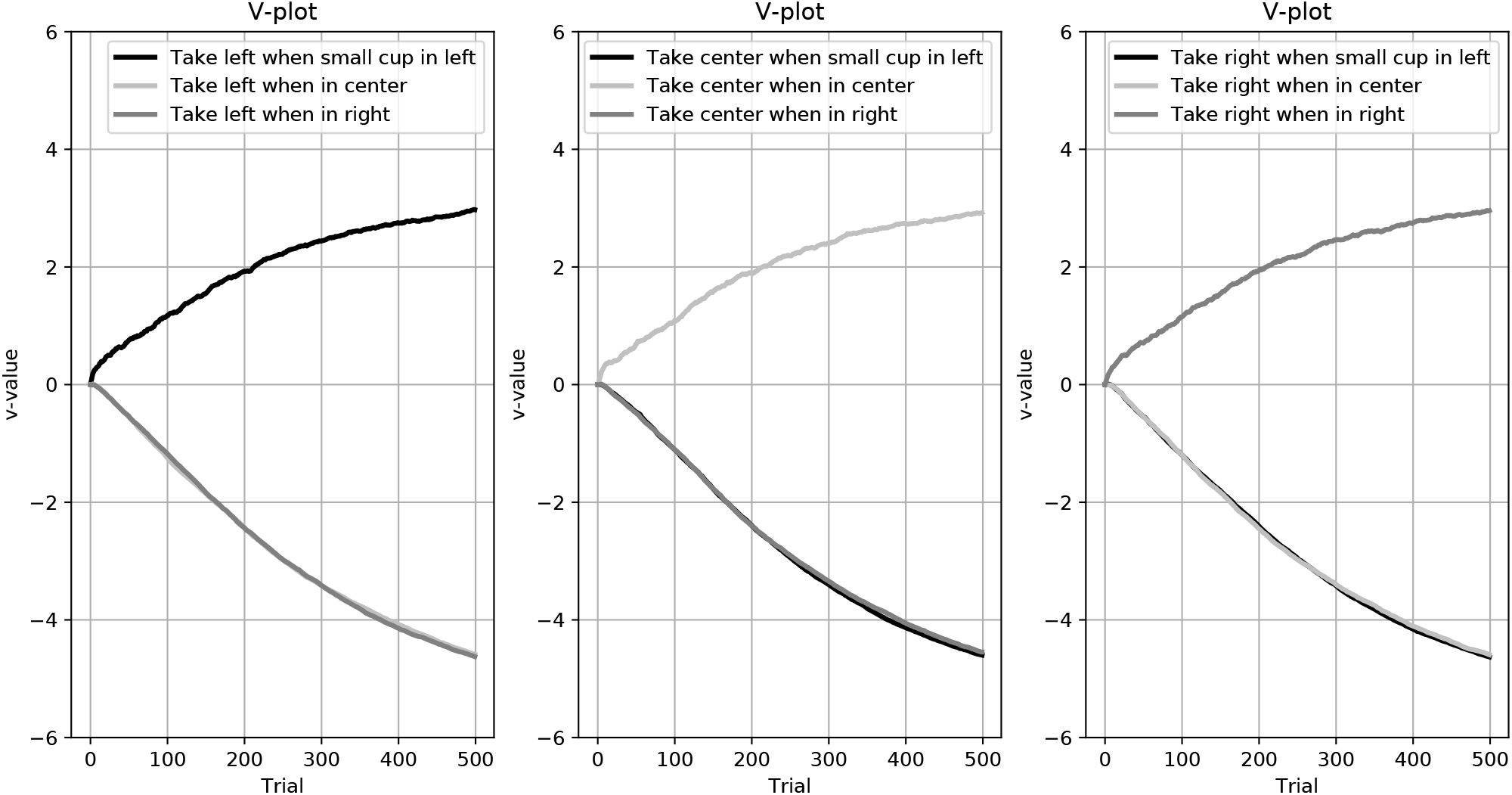
V-plot for the object permanence simulation.

#### 2.10 Numerosity

**Figure 10:**
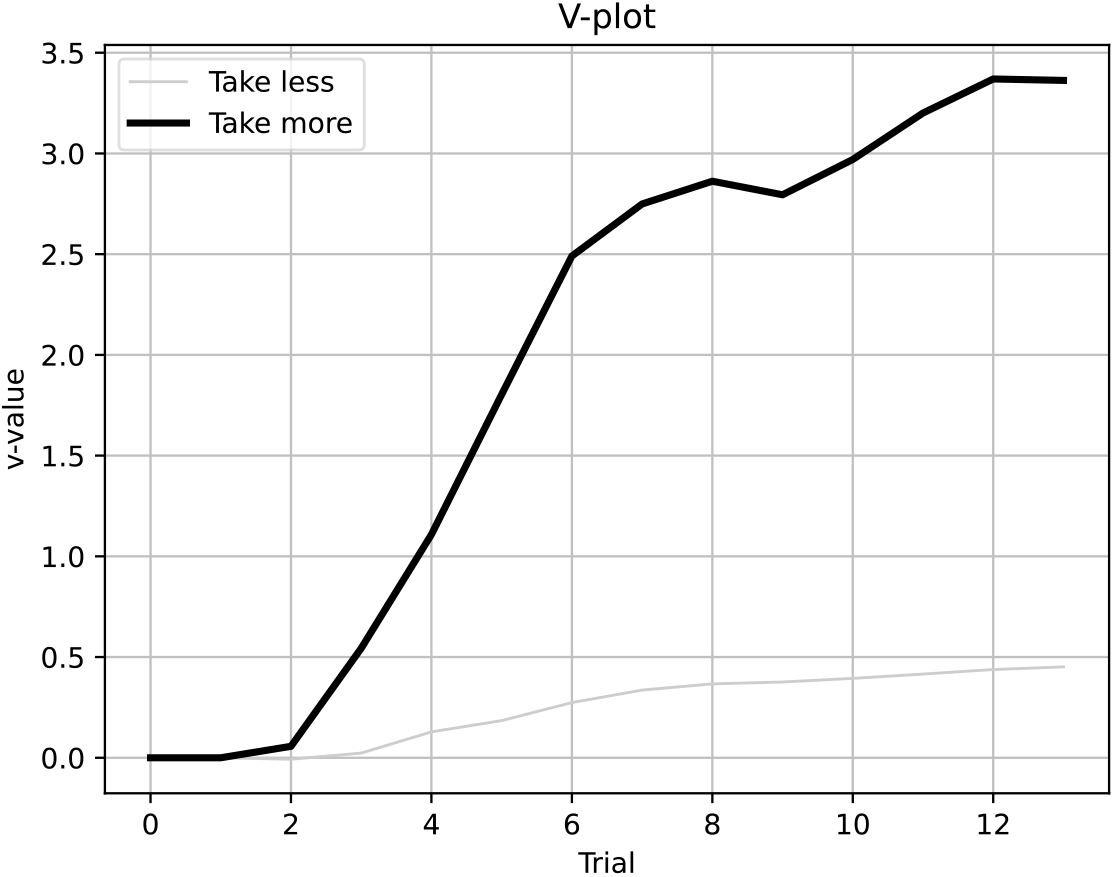
V-plot for the numerosity simulation for *s*(less) →*b*(take less) and *s*(more) →*b*(take more).

#### 2.11 Tool use

**Figure 11:**
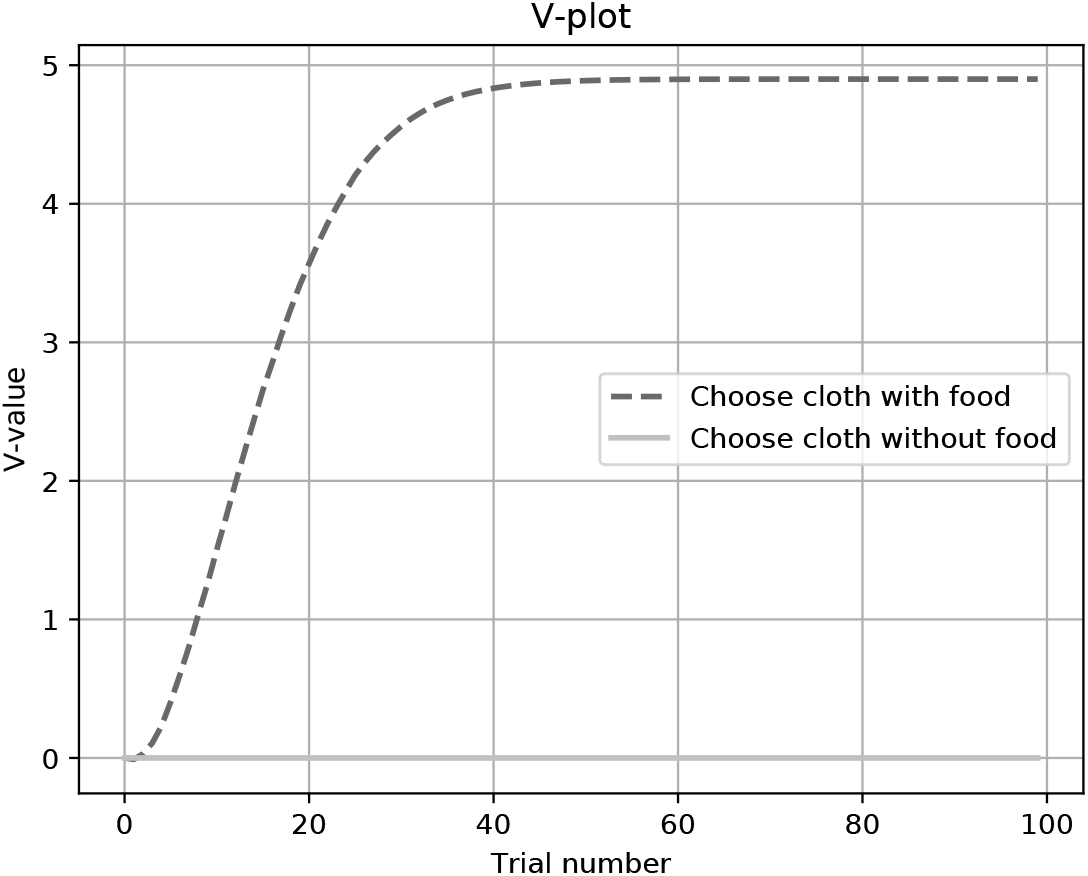
V-plot for the tool use simulation.

## Footnotes

1 At the extreme *β* = 0 behaviors are chosen randomly and memories from prior experiences do not affect decision making

## Notes

### Competing Interest Statement

The authors have declared no competing interest.

